# A dual target molecular MRI probe for noninvasive profiling of pathologic alpha-synuclein and microgliosis in a mouse model of Parkinson’s disease

**DOI:** 10.1101/2023.07.25.550555

**Authors:** Xianwei Sun, Andrew Badachhape, Jeannie Chin, Ananth Annapragada, Eric Tanifum

## Abstract

Parkinson’s disease is characterized progressive deposition of pathologic alpha-synuclein (α-syn) aggregates, neuroinflammation, and death of dopaminergic neurons in the substantia nigra projecting to the striatum. Noninvasive *in vivo* profiling of α-syn aggregate accumulation and microgliosis by molecular imaging can provide insights on the underlying mechanisms of disease progression, facilitating the development of effective treatment. However, no classical imaging methods have been successful, despite several attempts. We demonstrate a novel method to noninvasive *in vivo* profiling of pathologic α-syn in combination with microgliosis using molecular magnetic resonance imaging (MRI), by targeting oligomeric α-syn in cerebrospinal fluid with nano scavengers (**T**) bearing a T1-relaxive Gd(III) payload. In this proof-of-concept report we demonstrate, *in vitro,* that microglia and neuroblastoma cell lines internalize cross-linked **T**/oligomeric α-syn agglomerates. Delayed *in vivo* T1-weighted MRI scans following intravenous administration in the M83 α-syn transgenic mouse line show statistically significant T1 signal enhancement in test mice versus controls. The *in vivo* data was validated by *ex-vivo* immunohistochemical analysis which showed a strong correlation between *in vivo* MRI signal enhancement, Lewy pathology distribution, and microglia activity in the treated brain tissue. Furthermore, neuronal, and microglial cells in brain tissue from treated mice displayed strong cytosolic signal originating from **T**, confirming *in vivo* cell uptake of the nano scavengers.

## Introduction

Parkinson’s disease (PD) is the leading movement disorder and the second most common neurodegenerative disorder. Current statistics also show that it is the fastest growing neurological disease worldwide, with a 95% increase in incidence from 1990-2019, a growth rate that surpasses that of Alzheimer’s disease (AD)^1^. In the United States alone, an estimated 1.5 million individuals suffer from the disease, with 50 to 60 thousand new cases each year^2^. Across the globe, an estimated 6.2 million individuals suffer from PD, a number projected to double by 2040^1^. Despite this pressing need, the pathophysiological mechanisms underlying disease initiation and progression which can lead to the development of disease-modifying therapies remain elusive^1–3^. In fact, some experts now believe PD should be classified as a pandemic to heighten activism, focused planning, resource mobilization and novel approaches for detection, treatment, and management of the disease^3,4^.

The pathogenesis of PD mirrors that of related neurological disorders that are characterized by pathological deposits of misfolded protein aggregates, neuronal death, chronic neuroinflammation, and compromise of the blood-brain barrier (BBB)^5^. In the movement disorders generally referred to as synucleinopathies (which include PD, dementia with Lewy bodies, and multiple system atrophy) the pathological protein aggregates observed in postmortem brain tissue sections appear as cytosolic inclusions in neurons and glia, and are referred to as Lewy bodies and Lewy neurites^6^. Studies on regional distribution in postmortem PD brain tissue suggest that Lewy pathology originates in the olfactory system, peripheral autonomic nervous system, and dorsal motor nucleus of the glossopharyngeal and vagal nerves, and progressively spreads to other areas of the central nervous system (CNS). In addition, Lewy pathology in the medulla oblongata, pontine tegmentum, and anterior olfactory bulb pre-date clinical manifestations of PD-related motor symptoms. These symptoms first appear when the pathology has spread to the substantia nigra and other foci within the basal portions of the mid-and forebrain^7–9^.

The dominant protein component in Lewy pathology consists of α-synuclein (α-syn) aggregates. Under normal physiological conditions, α-syn adopts a helical conformation but under pathological conditions it misfolds and aggregates as β-sheets^10^. The prevailing theory is that disease propagation proceeds by cell-to-cell transmission of α-syn aggregates, mediated by reactive gliosis^11–14^. Total α-syn levels in the cerebrospinal fluid (CSF) and blood may not vary significantly between normal and PD patients, but there is strong evidence that the fraction of oligomeric aggregates corresponds with disease severity and plays a significant role in disease pathogenesis^15–19^. In normal individuals, α-syn is expressed predominantly in neurons, but in the disease brain, activated glia may internalize pathological aggregates via receptor-mediated and phagocytotic pathways^12,16^. Some evidence suggests that neuronal cells release small amounts of monomeric and oligomeric α-syn by exocytosis^12^ and oligomeric species have been shown to trigger reactive gliosis^15^. Genetically engineered mouse models of the disease, such as the A53T α-synuclein transgenic line M83, develop age-dependent regional distribution of intracytoplasmic neuronal α-syn inclusions that parallel disease onset and progression in humans^20,21^. Although these models do not perfectly match the exact human condition, they serve as excellent tools in the drug discovery process.

Technologies that can help in the detection and quantification of the pathologic variants of α-syn and reactive gliosis have great promise in early detection as well as the development of effective treatments of PD. While there has been considerable success in the development of protocols to detect and quantify oligomeric α-syn in CSF and blood samples with high accuracy^19^, noninvasive *in vivo* molecular imaging tool for reactive gliosis and pathologic α-syn in the brain, which can be more impactful in the development of disease-modifying therapies remain elusive. Most research on noninvasive imaging of α-syn aggregates in PD has focused on small organic molecule-based positron emission tomography (PET) or single-photon emission computed tomography (SPECT) tracers for Lewy pathology^22–24^. However, unlike PET tracers for extracellular amyloid-β (Aβ) plaques in AD, which have recently been approved by the FDA^25^, accessing Lewy pathology *in vivo* is challenging for several reasons. First, the pathology is intracellular and therefore more difficult to target with a PET tracer. Second, Lewy pathology in PD brains is far less abundant than Aβ in AD brains and tracers would need exceptionally high affinity to achieve results comparable to their Aβ counterparts. Moreover, in cases such as Parkinson’s disease dementia, Lewy pathology often co-occurs with other structurally similar amyloids such as Aβ and tau^26^. This means α-syn tracers must be highly selective; however, all tracers reported to date have some degree of affinity with these other amyloids^23^. These challenges call for new approaches to profile PD pathogenesis *in vivo*.

Over the past decade, we have demonstrated in several mouse models of neurodegeneration that liposomal nanoparticles administered intravenously cross the BBB into the CSF via the choroid plexus and microbleeds, carrying milligram quantities of imaging contrast payload per milliliter ^27–30^. We recently published that a variant of the probe designed to carry a hyper-relaxive Gd(III) MRI contrast payload, targeted to Aβ pathology, shows high efficacy in separating Aβ+ from Aβ-mice and is currently in clinical trials as the first MRI-based molecular imaging agent for Aβ in humans^29,30^. We also recently reported a series of indanones and indandiones as high-affinity and highly specific ligands for α-synuclein aggregates^31^. We hypothesize that a variant of the Aβ-targeted probe—one in which the targeting moiety is replaced by one of these novel α-syn ligands (**Figure 1A**)—can access oligomeric α-syn in the CSF and ISF and act as a scavenger, forming cross-linked agglomerates of α-syn aggregates-nanoparticles (**Figure 1B**). Such agglomerates can then be internalized by both neurons and activated glia cells via similar mechanisms (conventional endocytosis, receptor-mediated, and phagocytosis) as native oligomeric aggregates in the disease brain (**Figure 1C**). However, the agglomerates’ larger sizes will make them much better substrates (>0.5 microns)^32,33^ for accelerated phagocytic uptake by activated glia than the endogenous oligomeric species (∼200 nm), resulting in rapid accumulation of detectable levels of the contrast agent in disease brains. A nontargeted probe in the same brain environment will not form any agglomerates and therefore will not show significant accumulation. Similarly, a targeted probe in a brain without α-syn aggregates will also fail to show significant accumulation. As proof of concept, we prepared an α-syn-targeted formulation (**T**) composed of the lipids in Figure **1A**, with one of our indanone ligands^31^ as the targeting moiety, and a nontargeted control formulation (**N**). Using these two formulations and synthetic α-syn fibrils, we ran a series of *in vitro* experiments to verify agglomerate formation and cellular uptake of the ensuing agglomerates. Our data comprising dynamic light scattering (DLS) profiles, transmission electron microscopy (TEM), and confocal microscopy images demonstrates effective formation of cross-lined **T**/α-syn fibril agglomerates and accelerated uptake by neuronal and activated microglial cell lines, but not by an activated astroglia cell line. The *in vitro* data was validated *in vivo* in a pilot study in the A53T α-synuclein transgenic line M83 mouse model, including animals with intermediate and late-stage disease in which Lewy pathology is fully developed in all brain regions. Our *in vivo* MRI data shows statistically significant accumulation of the contrast agent in transgenic mice treated with **T** (**TgT**) compared to controls, including transgenic mice treated with **N** (**TgN**) and wildtype age-matched cohorts treated with **T (WtT)**. The *in vivo* observations were corroborated by *ex-vivo* immunohistochemical analysis that showed correlation between *in vivo* MRI signal enhancement and **T** distribution in brain tissue sections from treated mice, and internalization of the contrast agent by neurofilament-H and IBA-1 reactive cells.

**Figure 1.**
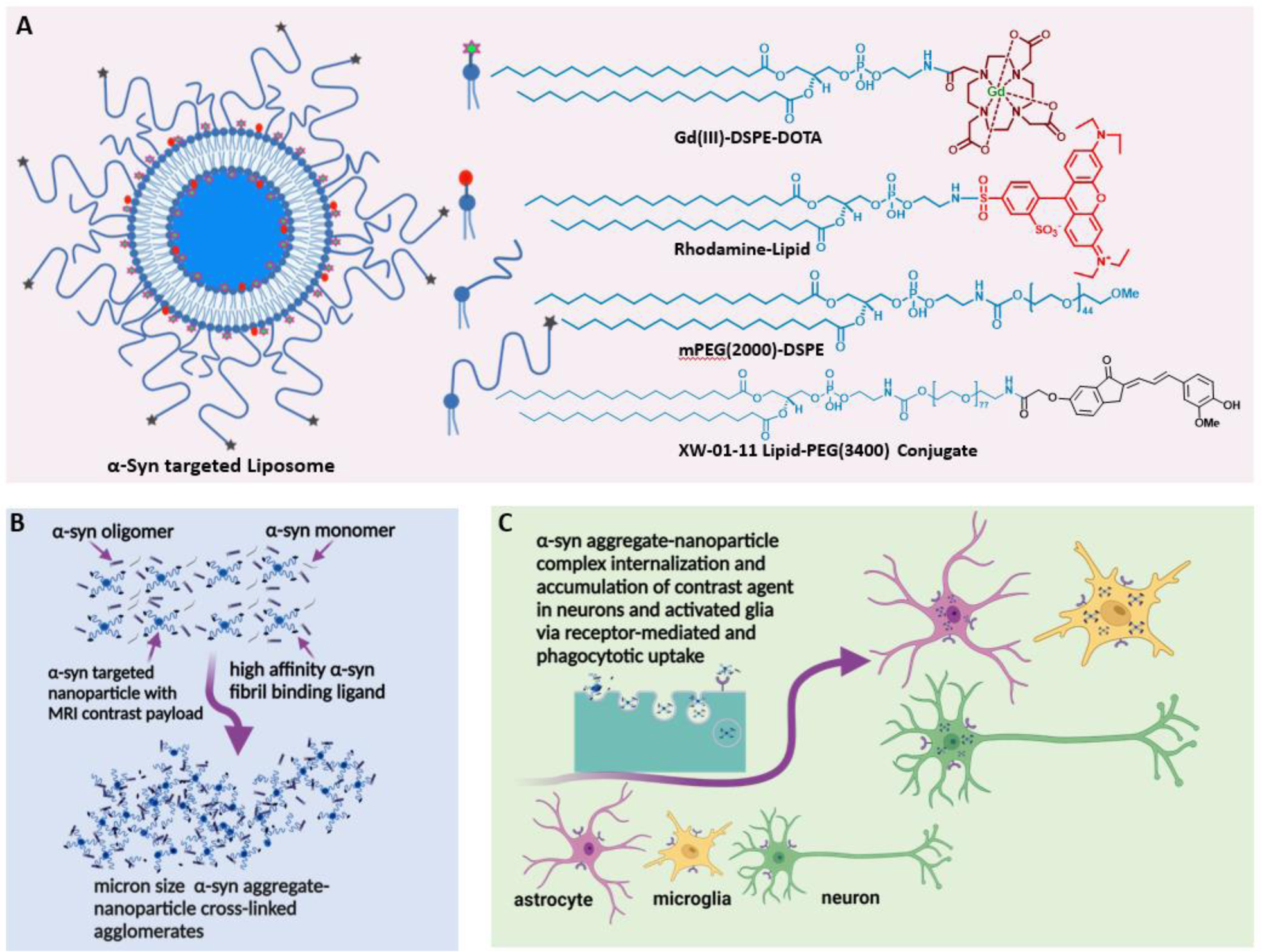
α-Synuclein nano scavenger hypothesis. **A**) Liposome nanoparticle labeled with oligomeric α-syn binding ligands (nano scavenger) and *in vivo* and *ex vivo* imaging contrast payloads; **B**) Nano scavengers cross-link oligomeric α-syn forming NS/oligomeric α-syn agglomerates in the CSF soup; **C**) Activated glia and neurons internalize agglomerates by conventional endocytosis, phagocytosis, and receptor-mediated uptake mechanism leading to momentary accumulation of detectable levels of contract agent in disease brains.

## II. Results

### II.1. Nano scavenger design and formulation

Design of the MRI-sensitive oligomeric α-syn-targeted nano scavenger was based on the highly T1-relaxive liposomal Gd(III), in which the macrocyclic Gd(III)-DOTA chelate is expressed on the lipid bilayer (Figure **1A**). We have previously demonstrated that this formulation generates Gd(III) solutions with about five times the T1 molar relaxivity of conventional Gd(III)-DOTA solutions^30^. Commercially available Rhodamine lipid was included in the formulation as a fluorescent reporter on probe in confocal microscopy imaging experiments. mPEG(2000)-DSPE was included to provide standard stealth properties. The targeting moiety, based on one of our previously reported α-syn aggregate-binding indanones (**XW-01-11**), was expressed on the surface of the liposome by a DSPE-PEG(3400)-Ligand conjugate. This conjugate was synthesized and characterized as shown in Supporting Information S1. Both **T** and **N** formulations were prepared using standard liposome formulation protocols. DLS measurements of **T** showed particles with a hydrodynamic diameter of 185.2 ± 5.3 nm and ICP-MS data showed a final lipid concentration of 88.4 ± 1.2 mM. The hydrodynamic diameter of **N** was 151.1 ± 2.2 nm and a lipid concentration of 84.0 ± 2.8 mM.

### II.2. In vitro verification and characterization of cross-linked oligomeric α-synuclein/nano scavenger agglomerate formation

To determine whether the **T** will form agglomerates upon exposure to oligomeric α-syn aggregates, we ran a series of *in vitro* experiments in which the **T** was exposed to synthetic α-syn fibrils under different conditions and then the ensuing products were characterized by DLS and TEM (Figure **2**). Synthetic α-syn fibrils were prepared as previously described^31^. A solution of fibrils (0.5 µg/mL) showed no DLS signal (Figure **2A.i**) while a solution of **N** at 10 µM lipid concentration showed particles with a hydrodynamic diameter of ∼150 nm (Figure **2A.ii**) and a solution of **T** (targeted formulation) at 10 µM lipid concentration showed particles with a hydrodynamic diameter of ∼185 nm (Figure **2A.iii**). We attribute the apparent difference in particle size between **N** and **T** to the PEG(3400) tether bearing the targeting ligands on the surface of **T** particles. Otherwise, the actual vesicle size is expected to be similar for both formulations since they were subjected to the same extrusion protocol. The DLS profile of solutions in which the lipid concentration of **N** is kept at 10 µM but with different fibril concentrations showed a very slight shift in the hydrodynamic diameter of particles in solution, as exemplified by Figure **2A.iv** (in which fibril concentration was titrated to 0.5 µg/mL). When fibrils (0.1 µg/mL final concentration) were added to a 10 µM lipid solution of **T**, two distinct particle populations were observed: one consisting of the original particles and a second population of larger particles, some with a diameter of up to a micron (Figure **2A.v**). An increase in the concentration of fibrils (as exemplified by the profile in Figure **2A.vi**, with a fibril concentration of 0.5 µg/mL), led to an increase in the population of the larger particles, suggesting agglomerate formation upon exposure of **T** to fibrils but not **N**. To ascertain whether agglomerate formation was due to **T** surface ligands binding to fibrils, blocking experiments were conducted in which **T** and fibrils were co-incubated with the free ligand at different concentrations. As exemplified by the DLS profiles in Figures **2A.vii** and **2A.viii**, respectively, co-incubation with 0.1 µM free ligand reduced the size of the larger particle population, and complete saturation of binding sites on the fibrils with 0.25 µM free ligand concentration in solution resulted in no agglomerate formation and a particle distribution like that observed after incubation of **N** with fibrils. Taken together, the foregoing data sets suggest effective *in vitro* agglomerate formation upon exposure of **T** to fibrils. A full panel of DLS profiles from Fibril dilution and blocking studies are shown in Figure **S1.1** in the Supporting Information section.

**Figure 2.**
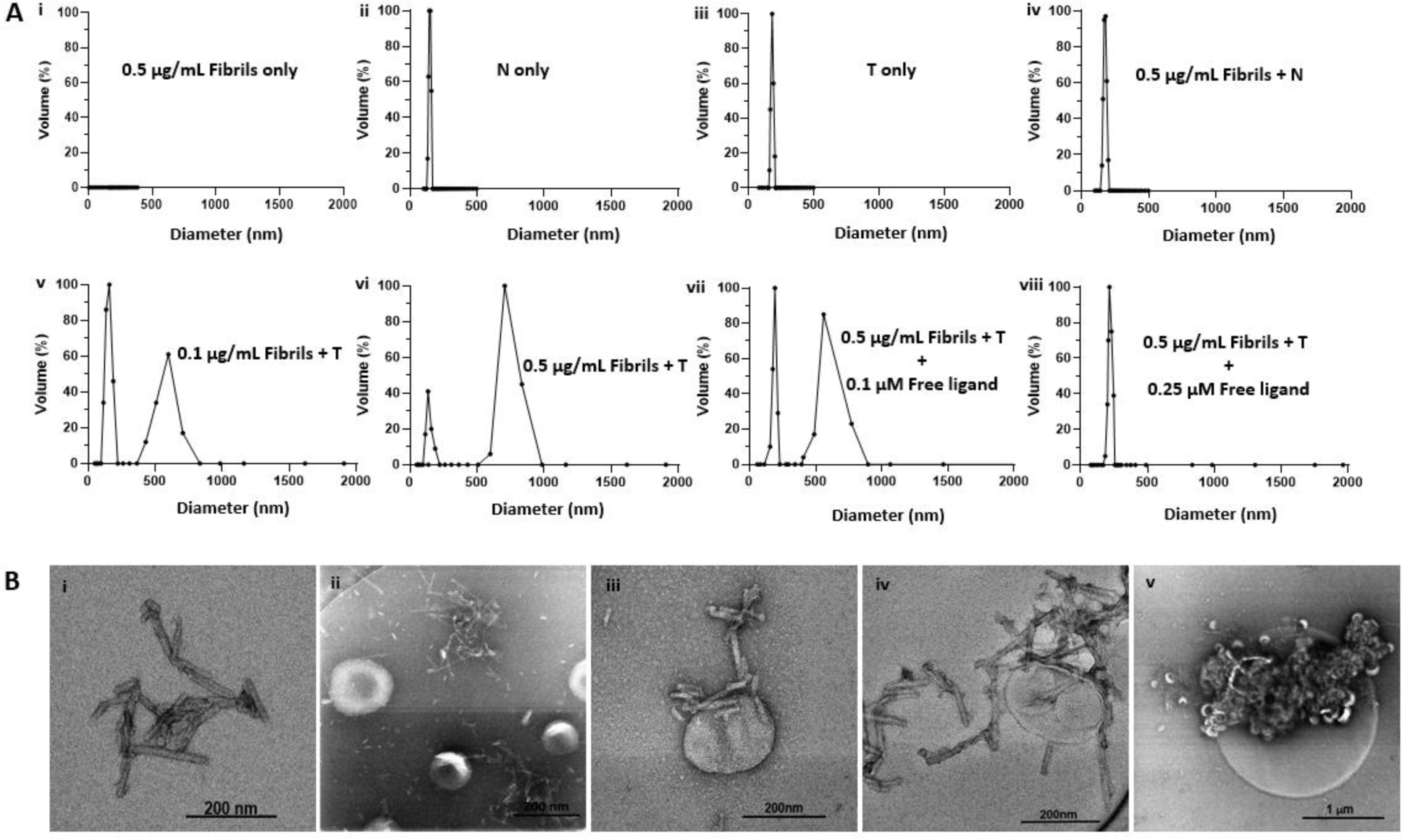
*In vitro* characterization of **T**/α-syn fibril agglomerate formation by DLS and TEM. All DLS experiments were run at 10 µM lipid concentration, for both **T** and **N**. **A**) DLS profiles of solutions of **T** or **N** at different fibril concentrations: **A.i**) 0.5 µg/mL solution of synthetic α-fibrils show no DLS signal; **A.ii**) Solution of **N** shows particle distribution of ∼150 nm; **A.iii**) Solution of **T** shows particle distribution of ∼185 nm; **A.iv**) 0.5 µg/mL solution of incubated with **N** shows no major change particle distribution; **A.v**) 0.1 µg/mL solution of fibrils incubated with **T** shows the original particle population with a hydrodynamic diameter of ∼185 nm and a new population with a hydrodynamic diameter of ∼600 nm, attributed to **T**/fibril agglomerate formation; **A.vi**) Increase in fibril concentration from 0.1 µg/mL to 0.5 µg/mL results in an increase in hydrodynamic of larger particles to ∼750 nm, attributed to increase in cross-linking at higher fibril concentration; **A.vii**) Maintaining fibril concentration at 0.5 µg/mL but introducing free fibril-binding ligand at 0.1 µM concentration results in an increase in the ∼180 nm volume fraction and a decrease in the ∼750 nm volume fraction; **A.viii**) Saturationof ligand binding sites on the fibrils with 0.25 µM free ligand results in no agglomerate formation; **B**) Negative stain transmission electron microscopy (TEM) images of α-syn fibrils exposed to **T**: **B.i**) TEM image of α-syn fibrils; **B.ii**) TEM image of **N** incubated with α-synuclein fibrils **B.iii**) TEM image of α-synuclein fibrils bound to a single **T** particle from a solution of **T** incubated with fibril; **B.iv**) TEM image of a nanometer size cluster of **T** bound to fibrils; **l**) TEM image of micron size **T**/fibrils agglomerate formed in a 0.5 µg/mL solution of synthetic α-fibrils incubated with **T**.

To further confirm the nature of **T**/fibril interactions, samples from solutions of fibrils and solutions of fibrils incubated with **T** were observed under transmission electron microscopy (Figure **2B**). Figure **2B.i** represents a sample TEM image of fibrils seen on grids prepared from solutions containing fibrils only while Figure **2B.ii** represent a sample image from a grid prepared from a solution of **N** incubated with fibrils. As seen in Figures **2B.iii** to **2B.v**, TEM images from grids prepared from samples solutions of **T** incubated with fibrils show a variety of agglomerate species. Figure **2B.iii** shows several fibrils bound to a single liposome, while Figure **2B.iv** shows a cluster of fibrils and **T** particles bound together, and Figure **2B.v** shows a micron-sized cross-linked **T**/fibrils agglomerate.

### II.3. In vitro accelerated cell uptake of agglomerates

The three key cellular species associated with the spread of α-syn pathology in PD have been identified as neurons, astrocytes, and microglia^12^. Neurons release small amounts of monomeric and oligomeric α-syn by exocytosis^12^ and express receptors involved in extracellular α-syn uptake^16^. Astrocytes undergo structural changes (i.e., are activated) upon exposure to oligomeric and fibrillar α-syn^34^, and it has been suggested they internalize aggregates through endocytosis^35^ and effectively degrade them via proteasomal and autophagic pathways^36^. Microglia become activated when exposed to soluble oligomeric α-syn or preformed fibrils^37,38^ and upregulate the expression of genes encoding receptors involved in extracellular α-syn uptake and degradation^16,38^. The primary uptake mechanisms are pattern recognition and phagocytosis but overloaded activated microglia have also been shown to transfer internalized α-syn cargo to neighboring naïve microglia via F-actin-dependent intracellular connections where it is rapidly degraded^39^. Another recent study demonstrated that microglia, aided by microglial Toll-like receptor 4 (TLR-4), engulf and α-syn into autophagosomes for degradation via selective autophagy^40^.

To verify these cells’ potential to internalize **T**/α-syn agglomerates as hypothesized, we ran *in vitro* cell experiments in which three different cell lines—SH-SY5Y (neuroblastoma cell line), T98G (astrocyte cell line), and HMC-3 (microglial cell line)—were exposed to preformed α-syn fibrils and **T** under various conditions. First, to ensure that the chosen glial cell lines became activated upon exposure to the fibrils, they were each incubated with fibrils for 24 hours and the incubation media assayed for cytokines. Controls included cells treated with liposaccharide (LPS) and untreated cells. **T** was also subjected to the same protocol to evaluate its inflammatory potential. The results (Figure **3A**) show that both HMC-3 and T98G cells exposed to fibrils demonstrate significant cytokine release relative to untreated controls and **T** alone. (Statistical analysis for this experiment was performed through one-way ANOVA with multiple pairwise comparisons between groups using Fisher’s least significant difference (LSD) procedure). Following this confirmation, we assessed the interaction of each of the three different cell lines upon exposure to **T** and fibrils. Controls included cells incubated with fibrils and **N**, and cells incubated with **T** alone. As exemplified by confocal microscopy images in Figure **3B**, HMC-3 cells incubated with fibrils and **T** showed unequivocal interaction with **T** (shown by the red color structures in the Rhodamine channel) within 1.5 hours of incubation, while cells incubated with either **T** only or **N** and fibrils showed no apparent interaction with the probe. Composite images generated by merging signal from nuclei stain (DAPI) and cytoskeleton signal (Phalloidin-488 nm) show that the red signal is cytosolic. Taken together with the control data, this observation suggests that the presence of fibrils is a requisite to **T** internalization. A summary of data from all three cell lines subjected to the same incubation protocol (Figure **3C**) showed that the astrocyte cell line does not avidly internalize the probe like its neuronal and microglial counterparts.

**Figure 3.**
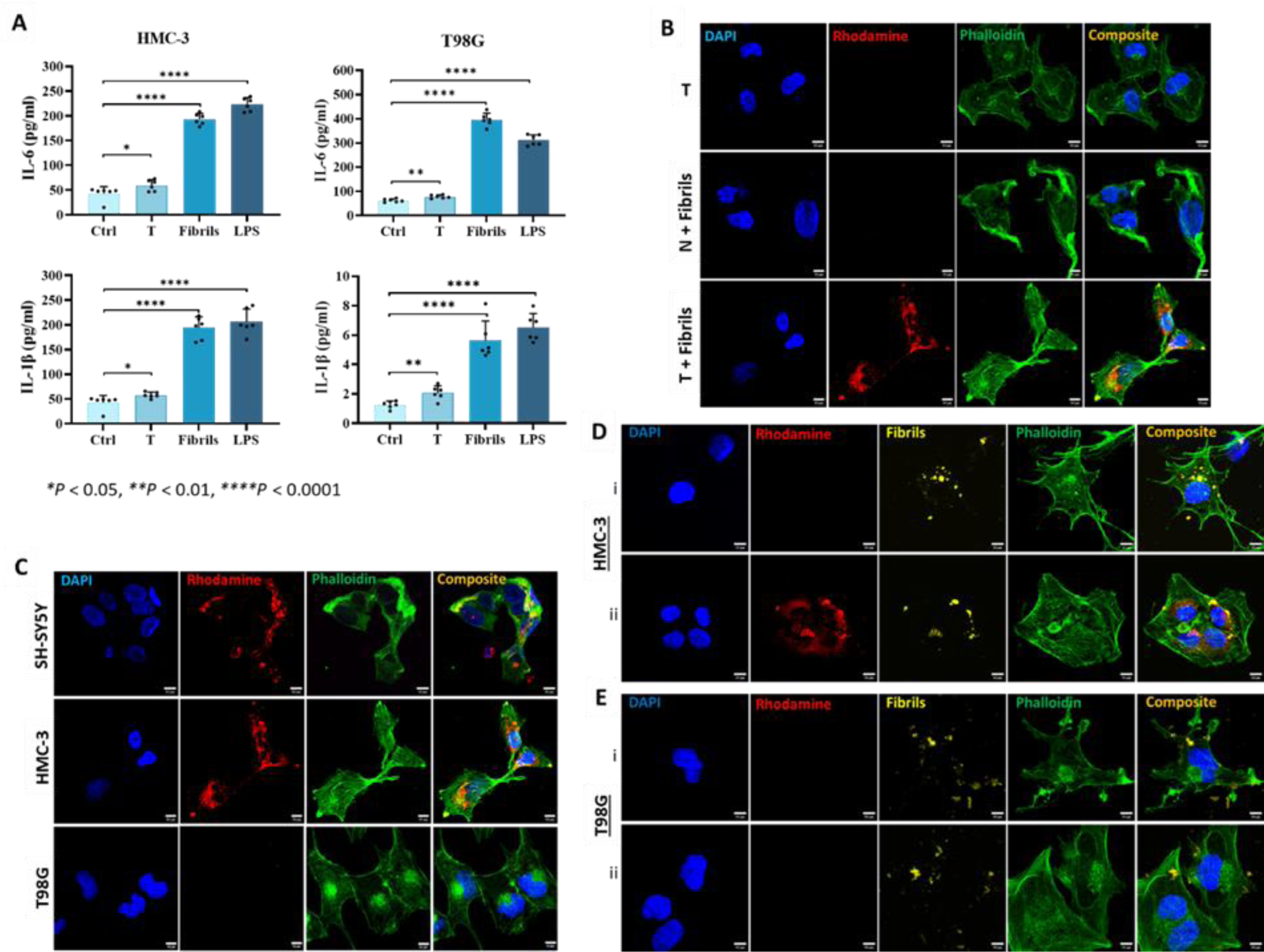
*In vitro* exposure of T to neuronal and microglia cell lines in the presence of α-fibrils results in accelerated internalization of T. **A**) Exposure of HMC-3 and T98G cells to α-syn fibrils (1.0 µM) results in the release of elevated levels of cytokines, including IL-6 and IL-1β, confirming activation; **B**) Confocal microscopy images of HMC-3 cells (microglia cell line) incubated with T only or N plus fibrils show no apparent cell interaction with the nano scavengers (Rhodamine, red fluorescence) within 1.5 h of incubation, while cells incubated with T and fibrils show a conspicuous red signal. Composite images of nuclei stain (DAPI, blue) and the cytoskeleton (Phalloidin-488 nm, green fluorescence) show that the Rhodamine signal is cytosolic; **C**) Summary of all three cell lines subjected to the same evaluation protocol suggests that only the neuronal and microglia cell lines avidly internalize the nano scavengers while the astrocyte cell line does not; **D.i**) HMC-3 cells incubated with labeled fibrils (Alexa Fluor^TM^-750 nm) show fibril internalization in the absence of T; **D.ii**) HMC-3 cells incubated with incubate with T and labeled fibrils show internalization of both species. More importantly, colocalization of the T and labeled fibril signal in the cytosol; **E.i**) T98G cells incubated with labeled fibrils do not show the avid internalization seen in their HMC-3 counterparts; **E.ii**) T98G cells incubated with T and labeled fibrils show neither avid uptake nor colocalization of signals from the two species in the cytosol. Statistics: One-Way ANOVA (Pairwise – Fisher’s LSD): *p<0.05,**p<0.01, ****p<0.0001. Scale bar = 20 µm.

To gain more insight into the role of fibrils in **T** interalization, fibrils were labeled with a fluourescent tag (Alexa Fluor^TM^-750 nm) and incubated with HMC-3 cells in the absence of **T**. As shown in Figure **3C.i**, within 1.5 hours of incubation, HMC-3 cells showed avid fibril internalization (yellow fluorescence) in the absence of **T.** When the HMC-3 cells were incubated with both **T** and labeled fibrils (Figure **3C.ii**), both species were internalized. More importantly, all the labeled fibril signal was colocalized with the strong and bright signal points originating from the Rhodamine channel (signal from **T**) in composite images (white arrows). This finding strongly suggests the formation and internalization of both large and small agglomerates in the process. The bright signal spots are attributed to agglomerates with a larger hydrodynamic diameter while the more diffuse signal points are attributed to smaller agglomerate species resulting from encounters between **T** and fibrils in the incubation mixture. Unlike their HMC-3 counterparts, the T98G cells (Figure **3D**) neither showed avid internalization of the fibrils by themselves (Figure **3D.i**) nor when co-incubated with **T** (Figure **3D.ii**). Overall, this data corroborates the observation above that **T** is internalized due to complexation with fibrils and that the astrocyte cell line does not internalize fibrils or **T**/α-syn fibrils agglomerates as avidly as the neuronal and microglial cell lines.

### II.4. In vivo molecular MRI evaluation of T

The A53T α-synuclein transgenic line M83 model, commonly referred to as M83 transgenic mice, expresses mutant human A53T alpha-synuclein under the direction of the mouse prion protein promoter. Some mice homozygous for the transgenic insert develop a progressively severe motor phenotype at eight months of age, but on average, the phenotype fully manifests at 14-15 months of age. Along with the motor impairment phenotype, they also develop age-dependent intracytoplasmic alpha-synuclein aggregate inclusions. Immunohistochemistry analysis of mutant mice between eight to 12 months of age reveal widely distributed alpha-synuclein inclusions, with dense accumulation in the spinal cord, brainstem, cerebellum, and thalamus. Pathology in older mice may also spread to other parts of the CNS, paralleling features of the disease in humans. For this proof-of-concept study, we chose two different age groups of transgenic mice in which the pathology is known to be fully established: 13 to15-month-old mice, representing intermediate stages of the disease, and 16+ month-old mice, representing late stages of the disease.

#### II.4.1. In vivo pharmacokinetics and biodistribution of nano scavengers

All our *in vivo* MRI experiments were conducted in a 1T permanent magnet MRI system. To confirm that our **T** exhibits a similar *in vivo* profile as the classical liposome formulation, 10 to12-week-old C57BL/6 mice were administered the agent by tail vein injection. T1 maps were collected around the brain, inferior vena cava (IVC), kidneys, liver, and spleen at different time points over fourteen days following intravenous administration. By day 14, signal in the blood and other organs had returned to baseline levels except in the liver and spleen. Any residual G(III) in these organs was further assessed over a 60-day period using ICP-MS. Data from the T1 maps and ICP-MS analysis is presented in Supplementary Information Figure **S1.2**. Taken together, the data suggests that the formulation has a systemic circulation half-life of 20.2 ± 0.6 hours and is cleared via the monocyte phagocyte system pathway, consistent with reported results.

#### II.4.2. Noninvasive interrogation of pathologic α-syn accumulation and reactive gliosis

Mice were subjected to pre-contrast scans using a T1-weighted 2D spin-echo (T1w-SE) sequence with 160 µm in-plane resolution and 1.2 mm slice thickness to obtain baseline images. Following these scans, the agent was administered via tail-vein injection (dose 0.1 mmol Gd(III)/kg) and the mice were returned to their cages. The test group within each age group consisted of six transgenic mice injected with the targeted formulation (**TgT**), while controls consisted of six transgenic mice injected with the nontargeted formulation (**TgN**) and six wildtype mice injected with the targeted formulation (**WtT**). At four days post-contrast administration, when all unbound contrast had cleared the from the blood pool, the animals were re-imaged using the same scan parameters as in the pre-scans to determine accumulation of contrast in the brain. Image analyses and signal quantification were performed with Osirix software (Pixmeo, Geneva, Switzerland). For quantitative signal analysis, signal from ∼1.2 mm thick coronal slices at three different positions around Bregma: 3.92 mm, Bregma: 0, and Bregma: −6.64 mm to represent signal in the olfactory bulb, parietal-temporal lobe, and brain stem, respectively. Signal beyond Bregma: −6.64 mm showed some inconsistencies attributed to magnetic field inhomogeneity at this position in the head coil of the magnet. As shown in the pseudo-colored images from a 19-month-old mouse (Figure **4A**), signal (pre-and post-contrast scans) from each brain region was windowed similarly for both the test and control groups (**TgT**, **TgN**, and **WtT**). Visual observation suggests higher signal intensity (green color) in the post-contrast images of **TgT** compared to the **TgN** and **WtT**. Signal change (%) between pre-contrast and delayed post-contrast images for each brain region was quantified by integration of signal in regions of interest shown by the white dotted lines. Box and whisker plots of the signal change in test animals versus controls (Figure 4B) show **TgT** mice illustrate statistically significant signal enhancement between post-contrast T1w-MRI and pre-contrast MRI relative to their **WtT** or **TgN** counterparts. Statistical analysis for in vivo experimental data was performed through Kruskal-Wallis method with multiple pairwise comparisons using the Bonferroni method. It should be noted that strong regional differences were seen in the olfactory bulb and parietal-temporal lobe, while the brainstem regions showed greater variance. Despite this variance, all three regions demonstrated strong *in vivo* MRI signal resulting from retention of the Gd(III) loaded targeted agents. Similar signal enhancement profile was observed in the brains of 13-to 15-month-old **TgT** cohorts (Supplementary data **S1.3**).

**Figure 4.**
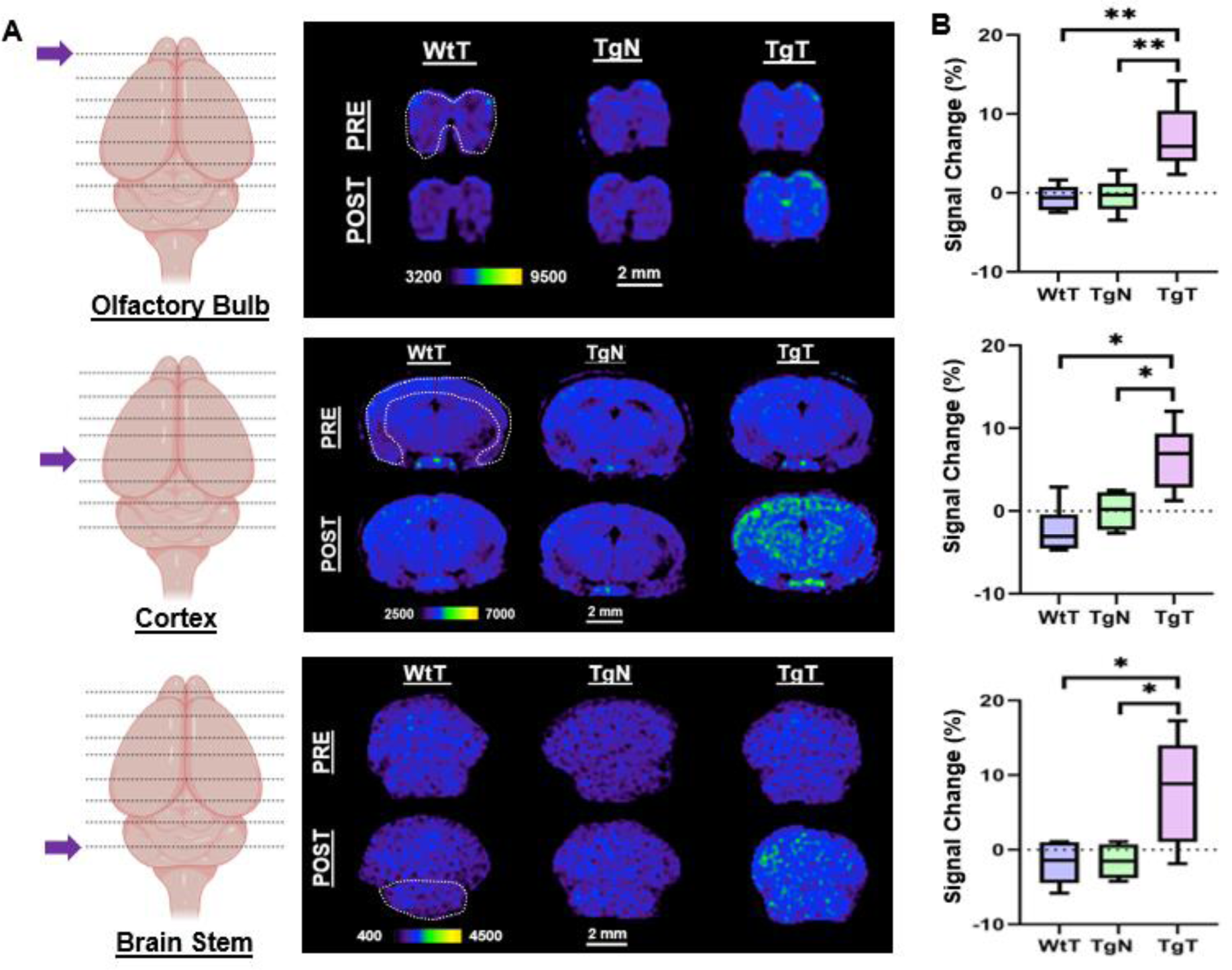
*In vivo* T1-weighted MRI pre-and post-contrast administration demonstrate statistically significant signal enhancement in M83 α-syn transgenic mice treated with **T** against controls. **A**) A comparison of 1.2 mm thick slices from T1-weighted spin-echo (T1w-SE) images at 4 days post nano scavenger administration around the olfactory bulb, cortex, and brainstem of 16 + month old **TgT** (N=6) against **TgN** (N=6), and **WtT** (N = 6) mice demonstrate signal enhancement in all the brain regions in TgT mice compared the **TgN** and **WtT** cohorts; **B**) Box and whisker plots of percent signal enhancement in all three brain regions show statistically significant mean enhancements in **TgT** against **TgN** and **WtT**. Statistics: Kruskal-Wallis (Pairwise – Bonferroni: * - p<0.05, ** p<0.005.

#### II.4.3. Ex-vivo immunohistochemical analysis

To confirm that the observed *in vivo* MRI signal enhancement in **TgT** mice was due to an interplay between α-syn aggregates and cellular participants, brain tissue from all the mice were assessed with immunohistochemical staining and confocal microscopy imaging following MR imaging. First, to confirm that the observed *in vivo* signal enhancement was due to the presence of the Rhodamine-labeled **T** in the tissue, signal from the Rhodamine channel in confocal microscopy images of the **TgT** tissue were compared against the **TgN** and **WtT** controls. As shown in Figure **5**, **TgT** brains showed fluorescence in the Rhodamine channel, suggesting the presence of **T** in the tissue. On closer examination, the fluorescence appears to emanate from two different sources: one that is very bright (white arrows), suggesting a more compact structure with high concentration of the fluorophore, and another which appears a little blurred (yellow arrows), suggesting somewhat more diffuse structures. A merge of the Rhodamine signal with a DAPI-stained nuclei image of the tissue showed all the Rhodamine signal in very close proximity to nuclei, suggesting the signal could be cytosolic. Very little to no Rhodamine signal was seen in images from the control tissue, consistent with the little to no positive signal enhancement in the *in vivo* MRI and corroborating the fact that the signal enhancement was due to **T** accumulation in the tissue.

**Figure 5.**
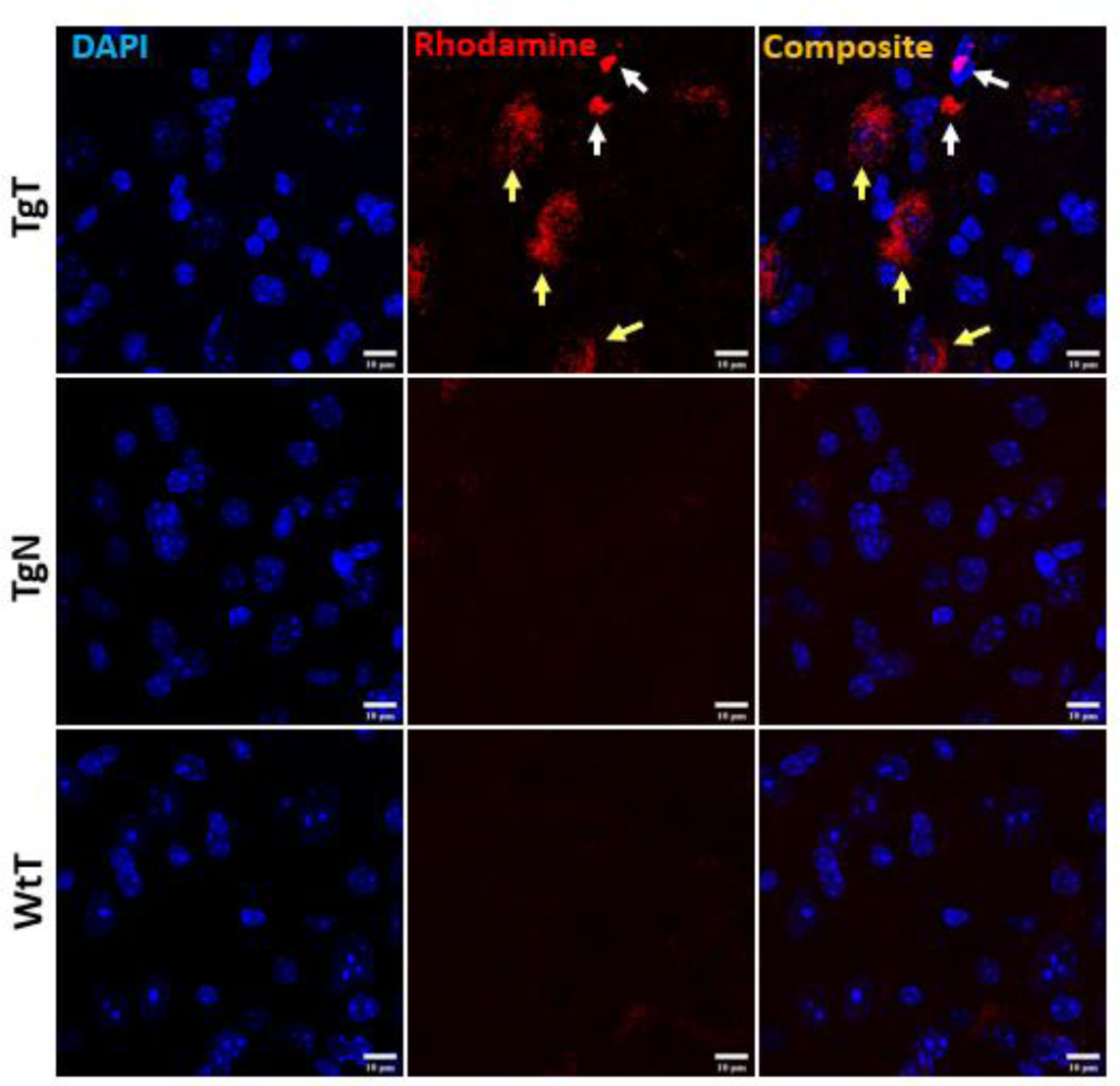
Confocal microscopy imaging demonstrated a correlation between *in vivo* MRI signal enhancement and accumulation of nano scavengers in brain tissue. Tissue sections from **TgT** mice show strong fluorescence in the Rhodamine channel compared to **TgN** and **WtT** controls. Closer examination revealed that part of the red fluorescence emanated from compact structures (white arrows) and the other part from diffuse structures (yellow arrows). Scale bar = 10 µm.

##### II.4.3.1 Correlation between in vivo MRI signal enhancement, nano scavenger signal, and phospho-S129-α-synuclein (pS129-α-syn) immunoreactivity

To assess the relationship between the observed *in vivo* MRI signal enhancement (shown above to correlate with **T** signal) and Lewy pathology, the imaged brain tissue sections were further subjected to immunohistochemistry with an antibody to phospho-S129-α-synuclein (pS129-α-syn), an established marker of pathological α-syn^41,42^. All brain regions that displayed *in vivo* MRI signal enhancement (olfactory bulb, parietal-temporal lobe, and brain stem) also showed Lewy pathology in association with **T** signal. As exemplified by images of cortical tissue sections from a 19-month-old **TgT**, (Figure **6A**), the tissue sections showed signal from **T** while strong anti-pS129-α-syn staining (750 nm) in cell bodies (Lewy bodies) was clearly visible throughout the entorhinal cortex (green fluorescence). Lewy pathology in the cortex and olfactory bulb is not widely reported in this model, likely because most studies have used mice younger (2 to12 months old) than the those in this study.

**Figure 6.**
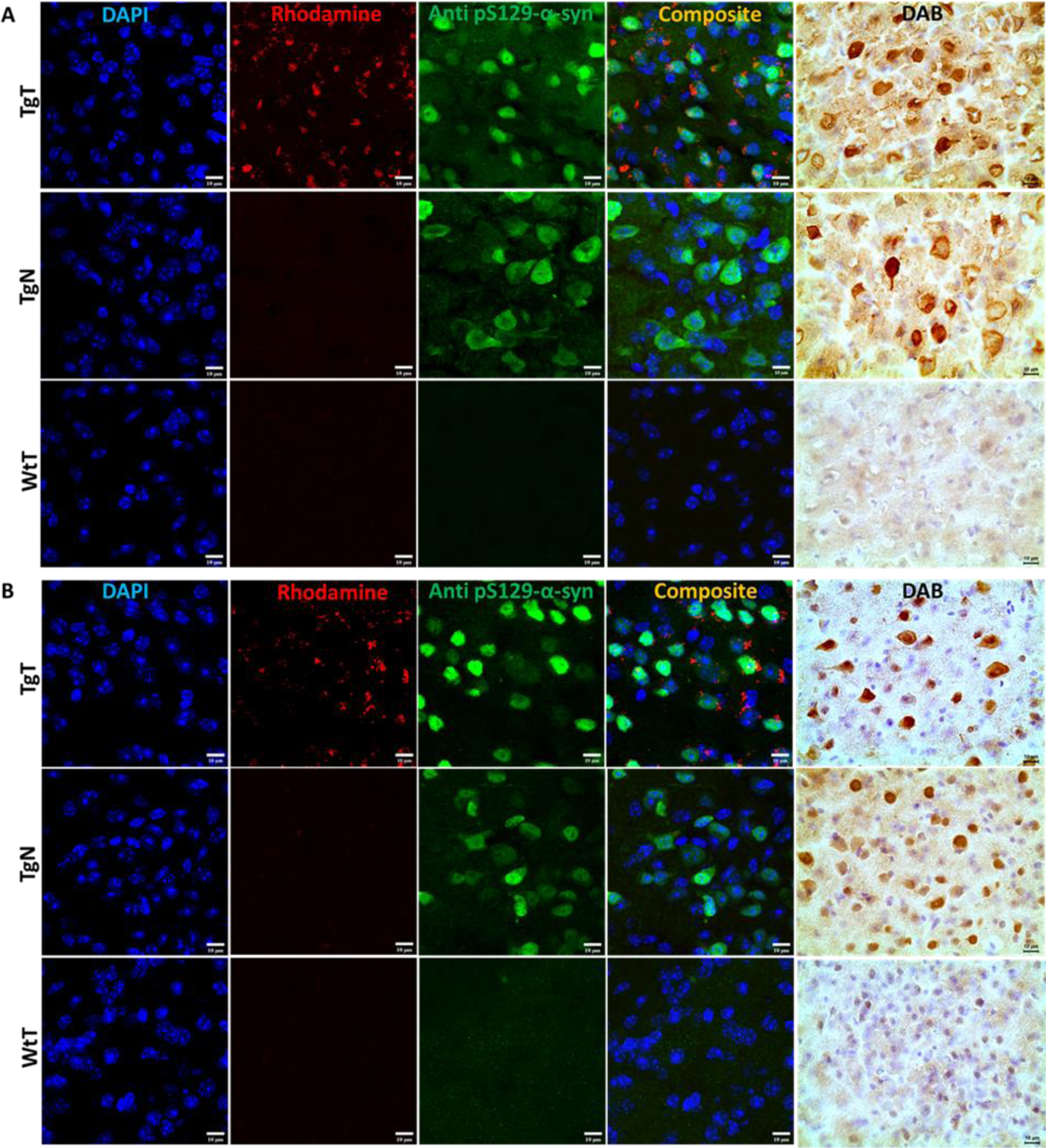
Immunohistochemical analysis demonstrates a correlation between *in vivo* MRI signal enhancement and nano scavenger location with Lewy pathology distribution. **A**) Anti-pSer129-α-syn stained cortical tissue sections from a 19-month-old **TgT** mouse show Rhodamine fluorescence in the entorhinal cortex compared with little to no signal in **TgN** and **WtT** controls. Secondary antibody fluorescence (750 nm) with characteristic cell body pS129-α-syn pathologic structures is observed in **TgT** and **TgN** tissue but not the **WtT** tissue sections. Contiguous slides subjected to the DAB protocol using pS129-α-syn (tailored for high specificity) confirm the observed structures as Lewy pathology; B) 13-15 month old cohorts display similar patterns of Lewy pathology, albeit less dense

To confirm that the observed green fluorescence was indeed Lewy pathology, contiguous tissue sections were stained with a pS129-α-syn DAB staining protocol which can be tailored to stain pathological α-syn with high specificity and no cross-reactivity with other proteins^43^. Bright field images from the DAB-stained tissue (brown fluorescence) show intense staining of the characteristic punctate structures of Lewy bodies on the tissue. Although **TgN** controls did not show any strong signal from the **T**, consistent with little to no *in vivo* MRI signal enhancement, they show strong anti pS129-α-syn reactivity which is consistent with their genotype. Images from other brain regions and age groups are shown in Figures **S1.3** - **S1.5** in the Supporting Information section.

##### II.4.3.2 Correlation between in vivo MRI signal enhancement, nano scavenger signal, and ionized calcium-binding adaptor molecule 1 (IBA-1) immunoreactivity

IBA-1 is a well-established marker for microglia and macrophages and is upregulated during the activation of both cellular species. To assess the presence microglia and their possible contribution to the observed *in vivo* MRI signal enhancement, brain tissue sections stained to assess IBA-1 expression patterns. Images of stained tissue sections from 16 plus-month-old mice (Figure **7A**) showed strong IBA-1 expression (green color) in both **TgT** and **TgN** sections. However, the strong **T** signal in the **TgT** tissue was absent in the **TgN** tissue due to the non-retention of the nontargeted particles, consistent with the little to no *in vivo* MRI signal enhancement in this cohort. A composite image created by merging the nuclei-stained image (DAPI), **T**, and IBA-1 fluorescence signals showed colocalization of IBA-1 reactivity with the compact structures with high Rhodamine fluorescence intensity (white arrows). The composite image also showed that the compact and highly fluorescent structures are cytosolic, suggesting *in vivo* engulfment of pre-formed **T**/oligomeric α-syn agglomerates by microglia. Images from their 13 to 15-month-old counterparts (Figure **7B**) and sections from the olfactory bulb and brain stem regions (Supplementary Figure **S1.6** **- S1.11**) displayed similar Rhodamine signal retention patterns and correlation between Rhodamine signal and IBA-1 reactivity. Taken together, this data strongly suggests that **T** encounters and form agglomerates with oligomeric α-syn aggregates *in vivo*, and the ensuing compact structures are engulfed by activated microglia, resulting in the momentary accumulation of the contrast agent in this cell type as hypothesized. However, the highly fluorescent structures are just a fraction of the total signal since signal from the diffuse fluorescent structures does not appear to colocalize with IBA-1 reactivity. To account for this signal and the role of the other cellular participants, the tissue sections were further stained for Neurofilament H (NF-H) and glial fibrillary acidic protein (GFAP) to assess the roles of neurons and astrocytes, respectively.

**Figure 7.**
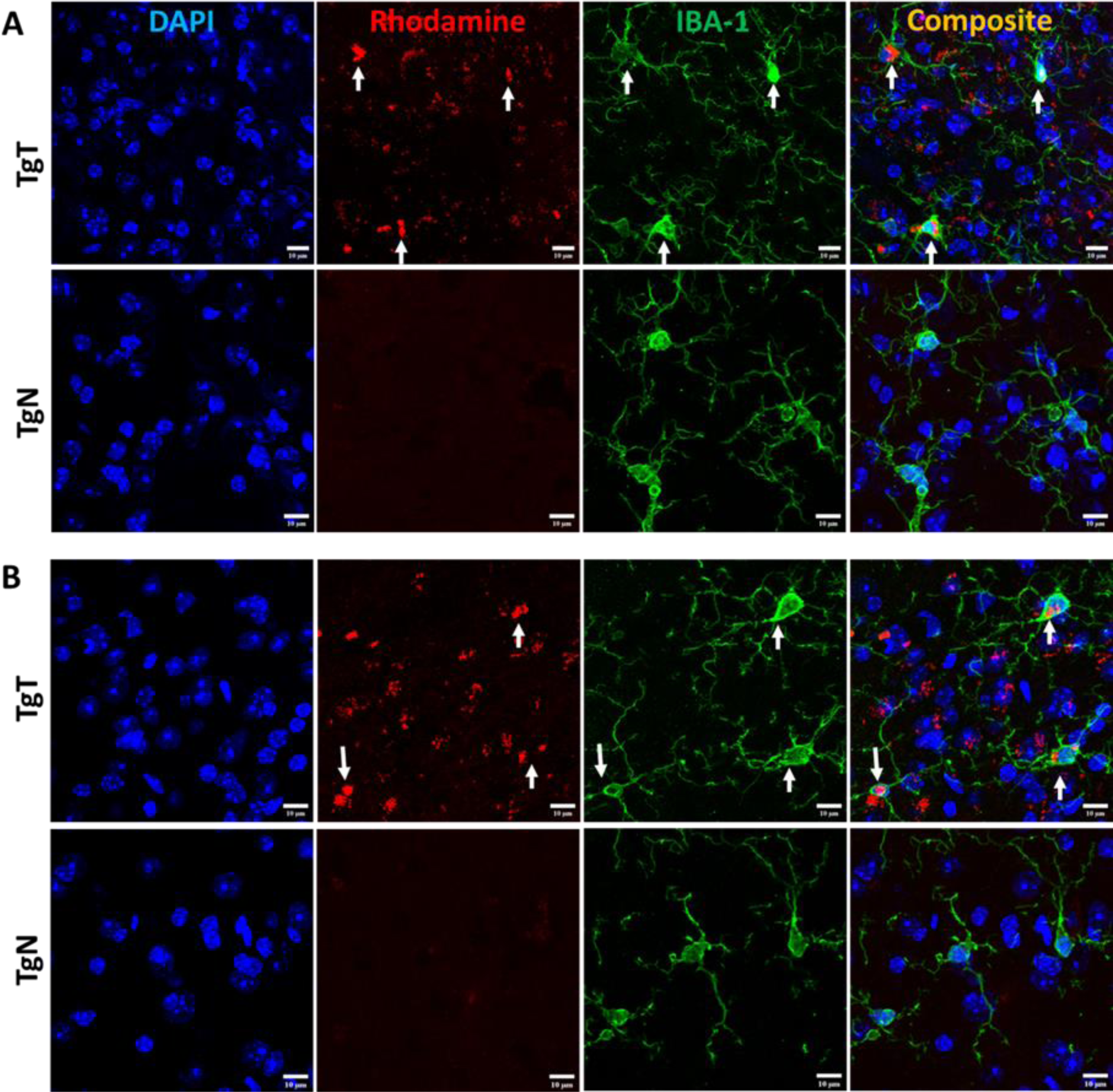
Microglia display *in vivo* uptake of nano scavenger species with apparent compact structural formation. **A**) Brain tissue sections from 19-month-old **TgT** (N=6) and **TgN** (N=6) mice show cell bodies and processes with strong IBA-1 reactivity (green color, 750 nm). **TgT** sections also show strong NS signal (Rhodamine, red color) while **TgN** cohorts show very faint to no signal. Composite images created by merging signal from nuclei stain (DAPI), the NS signal, and IBA-1 signals show that the strong compact NS signal is within the cytosol of the IBA-1 reactive cell bodies, **B**) Images from the younger cohorts demonstrate similar patterns.

##### II.4.3.3 Correlation between in vivo MRI signal enhancement, nano scavenger signal, and anti-neurofilament H (NF-H) immunoreactivity

Neurofilament H is one of three intermediate filament proteins (including NF-L, NF-M and NF-H) found specifically in neurons. Antibodies to NF-H are useful for identifying neuronal cells and their processes in tissue sections and tissue culture. To assess the location of neurons relative to the **T** signal in the tissue and their contribution to the observed *in vivo* MRI signal enhancement, brain tissue sections were stained for NF-H. As exemplified by images of stained tissue from a 19-month-old **TgT** (Figure **8A**), anti-NF-H reactivity (green color) was observed in tissue sections from all brain regions tested. Both cell bodies and axonal processes were also clearly visible. However, the cell bodies in the brain stem appeared to stain better than their counterparts in the cortical sections. The images also suggest the NF-H reactive cell bodies correlated mostly with the diffuse **T** signal (red color). A composite image generated by merging both signals with the nuclei stain (blue) shows that the diffuse **T** signal (yellow arrows) colocalizes with the anti-NF-H reactive cell bodies. Furthermore, the signal is cytosolic, suggesting uptake of **T**-labeled α-syn oligomers by neurons. Images from other brain regions and age groups are shown in Figures **S1.12** - **S1.13** in the Supporting Information section.

**Figure 8.**
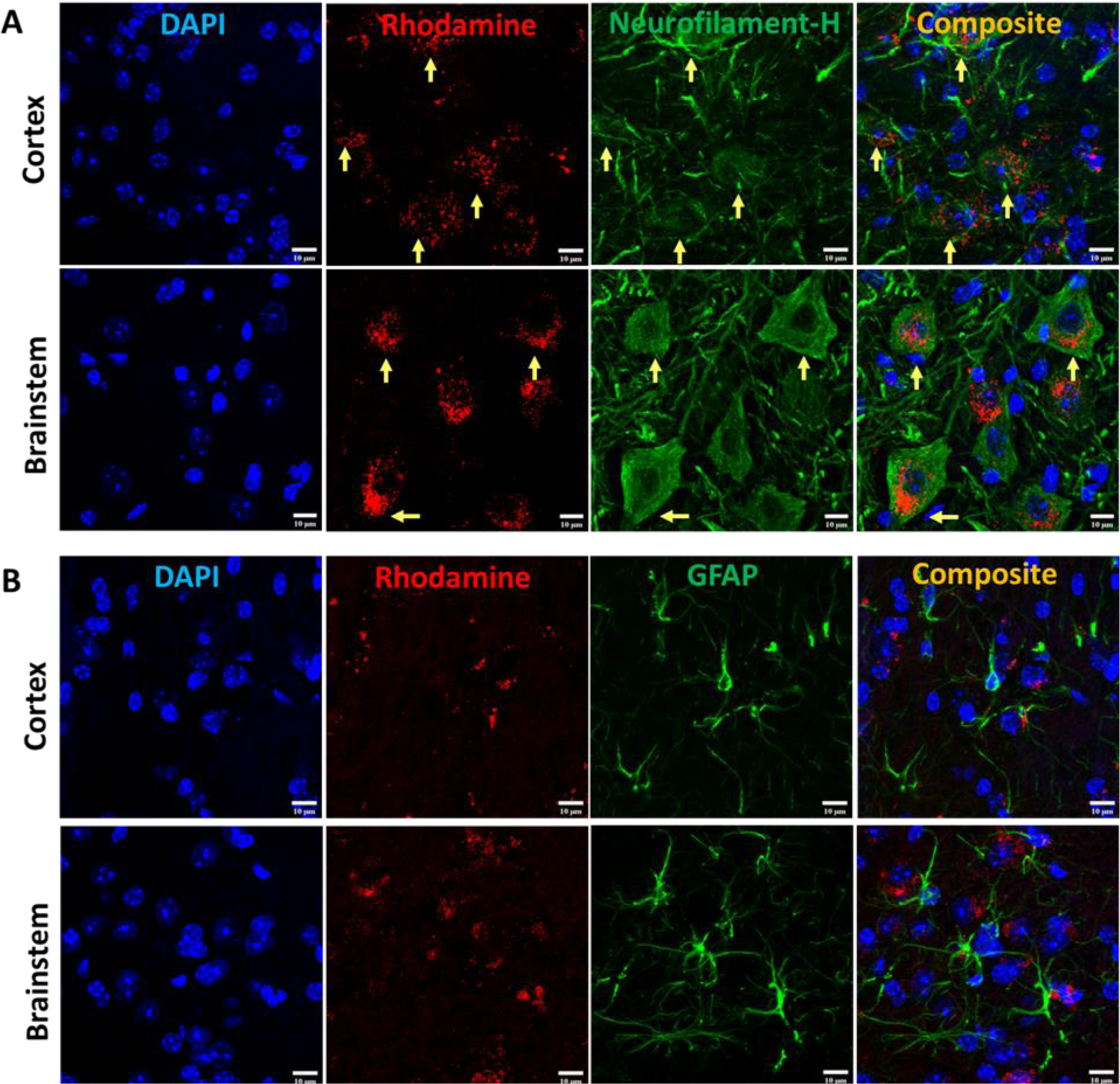
Neuronal cell bodies display *in vivo* uptake of nano scavenger species with apparent diffuse structural formation while astrocytes do not display convincing evidence of *in vivo* uptake. **A**) Anti NF-H stained brain tissue displays Rhodamine signal as well as cell bodies with strong anti-NF-H reactivity (green fluorescence, 750 nm) in both cortical and brainstem tissue sections, but the cell bodies appear to be more defined in the brainstem sections. A composite image of both signals show that the diffuse Rhodamine signal is cytosolic (yellow arrows). **B**) Anti GFAP stained sections show Rhodamine signal as well as strong GFAP reactivity, but composite images do not show conclusive evidence of internalization of **T** by GFAP reactive cells.

##### II.4.3.4 Correlation between in vivo MRI signal enhancement, nano scavenger signal, and glial fibrillary acidic protein (GFAP) immunoreactivity

GFAP is an intermediate filament protein principally found in astrocytes in the CNS, but it can also be found in neurons, hepatic stellate cells, kidney mesangial cells, pancreatic stellate cells, and Leydig cells. It acts as an intra-cellular structural component of the astrocytic cytoskeleton and expression the GFAP gene has attracted considerable attention because its onset is a marker of astrocyte development, and its upregulation is a marker of reactive astrogliosis. The GFAP antibody is well established as a useful marker of astrocyte reactivity. To understand **T** and astrocyte interactions *in vivo* and their relationship to the observed MRI signal enhancement, brain sections were immunoassayed with GFAP antibody. Images of stained tissue (Figure **8B**) showed strong anti GFAP reactivity with the characteristic asterisk structures of reactive astroglia in all brain sections of the **TgT** and **TgN** mice. This reactivity cooccurred with **T** signal (Rhodamine, red fluorescence), suggesting a correlation between astrocyte reactivity and the observed *in vivo* MRI signal enhancement. However, composite images generated by merging the two fluorescent images with the nuclei-stained image (DAPI) did not show the unequivocal cytosolic co-localization of the **T** signal in astrocytes as observed in microglia and neurons in any brain region. Images from other brain regions and age groups are shown in Figures **S1.14** - **S1.19** in the Supporting Information section.

## III. Discussion

This study strongly supports our nano scavenger/oligomeric α-syn/reactive gliosis/neuronal interplay hypothesis. Our *in vitro* data demonstrates that exposing **T** to α-syn fibrils resulted in the formation of agglomerates. The DLS data shows agglomerate species with hydrodynamic diameters ranging from 200 nm to >1 micron at fibril concentrations of 0.1 µg/mL. This particle size range was corroborated by TEM measurements, which showed different **T**/fibrils species within the same size range. Some literature reports suggest CSF concentration of oligomeric α-syn in PD patients as high as 170 ng/mL range^44,45^. The lack of agglomerates when **N** was exposed to fibrils at concentrations up to 0.5 µg/mL, the reduction in agglomerate size and volume, and no agglomerates formed in the saturated free ligand blocking experiments all demonstrate that agglomerate formation results from the specific binding of ligands on the surface of **T** to ligand binding sites on the fibrils.

Data from our *in vitro* cell uptake studies is equally in line with the hypothesis. Particles in the brain >0.5 microns in diameter are particularly suited for phagocytosis and microglia are the principal modulators of phagocytosis in the CNS^32,33^. While both microglial and neuroblastoma cell lines showed avid uptake of fibrils under all tested conditions, there was little evidence of uptake by the tested astrocyte cell line. More importantly, although the neuroblastoma and microglial cell lines internalized α-syn fibrils within 1.5 hours of exposure, **T** was not internalized within the same time frame unless in the presence of fibrils, while the **N** formulation, which lacks the targeting ligands, was not internalized in the presence of fibrils within the same time frame. These findings align with the assertion that oligomeric α-syn would aid in the internalization and accumulation of **T** bearing an imaging contrast payload in the cells.

More importantly, our *in vitro* data translated seamlessly into our *in vivo* results. As hypothesized, noninvasive *in vivo* brain MRIs showed statistically significant signal enhancement in **TgT** mice compared to controls due to accumulation of the contrast agent. Immunohistochemical analysis of the treated brain tissue revealed a strong correlation between regional distribution of the *in vivo* signal and Lewy pathology. Although there was no independent confirmation of the presence of oligomeric α-syn species in the tissue, we assume that the intracellular Lewy bodies and neurites are in flux with oligomeric species in the CSF and IF. IBA-1 reactivity revealed co-localization of microglia with the MRI signal. Cytosolic inclusions with strong **T** fluorescent signal were clearly visible within the microglia cell bodies. We speculate that this strong signal comes from larger pre-formed **T**/oligomeric α-syn agglomerate species engulfed by microglia via phagocytosis. However, our *in vitro* data and some literature reports suggest that microglia can engulf oligomeric α-syn into autophagosomes for degradation^40^ and one can expect that if fluorescently labeled, α-syn oligomers in autophagosomes could also appear as bright structures within the microglia cell bodies. Indeed, this is what we observed in the images of HMC-3 cells exposed to labeled α-syn fibrils. On the other hand, endogenous α-syn is not fluorescently labeled and the red fluorescence in the microglia in brain tissue sections of **TgT** mice can only originate from **T**. Taken together with our *in vitro* data, as well as our *in vivo* control data showing no signal enhancement in **TgN** mice, the cytosolic red fluorescence in the microglia is likely due to internalization of larger pre-formed **T**/oligomeric α-syn complex species in the CSF and IF by the cells. By this same reasoning, the diffuse cytosolic red fluorescence observed in the anti-NF-H labeled cell bodies was attributed to internalization of smaller pre-formed **T**/oligomeric α-syn species by neurons. This would most likely be via similar mechanisms as conventional endocytosis or receptor-mediated internalization of native oligomeric α-syn by neurons^12^. Surprisingly, despite several reports on astrocyte involvement in the spread of α-syn pathology^12,35^, we found no clear evidence of avid internalization of fibrils (with or without **T**) by the T98G cells *in vitro* or **T** signal in GFAP-expressing cells *in vivo*. However, these results are consistent with reports that microglia are the principal scavengers of extracellular oligomeric α-syn^12,17^.

## IV. Conclusion

This pilot study paves the way to a novel approach to simultaneously profile α-syn accumulation and microgliosis using *in vivo* molecular MRI in the M83 α-synuclein mouse model by a single injection. We hypothesized that: 1) nano scavengers labeled with oligomeric α-syn-binding ligands and bearing an imaging contrast payload would interact with soluble α-syn aggregates in the CSF and IF in a PD brain to form cross-linked **T**/oligomeric α-syn agglomerates of varying sizes; 2) the ensuing larger agglomerates would serve as good substrate for phagocytic uptake by reactive glia while the smaller species can be internalized by both neurons and glia via known receptor-mediated and other endocytosis pathways; and 3) the result would be momentary accumulation of the contrast agent at detectable levels in disease brains. Our *in vitro* data, assessed by DLS and TEM measurements, verifies the agglomerate formation portion of the hypothesis. We demonstrated that a spectrum of agglomerate species with sizes ranging from 200 nm to >1 µm in diameter are generated upon exposure of **T** to α-syn fibrils *in vitro*. Tested neuroblastoma and microglial cell lines showed avid internalization of fibrils with or without **T**, but avid rapid uptake of **T** is only possible in the presence of fibrils. The tested astrocyte cell line did not display unequivocal internalization of either fibrils or **T**, when used separately or in combination. *In vivo* MRI demonstrated that transgenic mice injected intravenously with the agent showed statistically significant brain signal enhancement due to retention of the agent, while *ex vivo* immunohistochemical analysis demonstrated that microglia and neurons were strongly implicated in retention of the agent in the brain. In addition, the *in vivo* brain MRI signal enhancement was shown to correlate with both Lewy pathology accumulation and microglial activity. To the best of our knowledge, no other report has demonstrated this type of interplay between nanoparticles and oligomeric α-syn, or any of the other amyloid aggregates with microglia and neurons.

Although the fate of internalized agglomerates in this study was not determined, a recent report demonstrated that microglia engulf α-syn aggregates into autophagosomes where they are digested. Our data demonstrating *in vitro,* and *in vivo* internalization of **T** is overwhelming, but we note that: the uptake mechanism is rather speculative; the fate of the internalized agent is not yet understood; and younger mice, which can offer greater insight regarding the disease’s early development and propagation, were not tested. We are currently investigating each of these areas and will report our findings in due course.

## Supporting Information

Detailed experimental protocols, supplementary images, NMR, and MALDI spectra.

## Acknowledgements

The authors acknowledge the writing support team at Texas Children’s Hospital, Houston, Texas for English language edits. This work was funded by the National Institutes of Health, USA (R21 AG067131 to Eric Tanifum.

## Ethical approval

Animal handling procedures were carried out following approved protocols by the Baylor College of Medicine Institutional Animal Care and Use Committee (IACUC).

## Data Availability Statement

The data that support the findings of this study are available from the corresponding author upon reasonable request.

## Supporting Information

### General information and methods

#### Materials

Unless otherwise noted, all reagents and solvents were obtained from commercial sources including, Sigma-Aldrich, TCI, and Acros Organics and used without further purification. Thin layer chromatography (TLC) was performed on silica gel 60 F254 plates from EMD Chemical Inc. and components were visualized by ultraviolet light (254 nm and) and/or phosphomolybdic acid, 20 wt% solution in ethanol. SiliFlash silica gel (230–400 mesh) was used for all column chromatography.

##### Measurements

Proton nuclear magnetic resonances (^1^H NMR) were recorded at 600 MHz or 500 MHz on Bruker 600 or 500 NMR spectrometers. Carbon nuclear magnetic resonances (^13^C NMR) were recorded at 75 MHz or 125 MHz on a Bruker 300 or 500 NMR spectrometers respectively. Chemical shifts are reported in parts per million (ppm) from an internal standard chloroform (7.26 ppm), or dimethylsulfoxide (2.50 ppm) for ^1^H NMR; and from an internal standard of either residual chloroform (77.00 ppm), or dimethylsulfoxide (39.52 ppm) for ^13^C NMR. NMR peak multiplicities are denoted as follows: s (singlet), d (doublet), t (triplet), q (quartet), dd (doublet of doublet), td (doublet of triplet), dt (triplet of doublet), and m (multiplet). Coupling constants (*J*) are given in hertz (Hz). High resolution mass spectra (HRMS) were obtained from The Ohio State University Mass Spectrometry and Proteomics Facility. The mean particle size of liposomes was determined using a Dynamic Light Scattering (DLS) instrument (Brookhaven Instruments Corp., Holtsville, NY, USA). Negative-stain Transmission Electron Microscopy images were captured from a JEOL 3200FSC (300kV) electron microscope. Magnetic Resonance Imaging (MRI) was performed on a 1 T permanent magnet scanner (M2 system, Aspect Imaging, Shoham, Israel). All confocal microscopy images were captured from an Olympus IX81 microscope and the images were processed and analyzed using Fiji-ImageJ software.

##### Chemical synthesis

The lipid conjugate bearing the targeting moiety, **DSPE-PEG(3400)-XW-01-11 Conjugate** was synthesized following synthetic route in Scheme **1**.

**Scheme S1.1.**
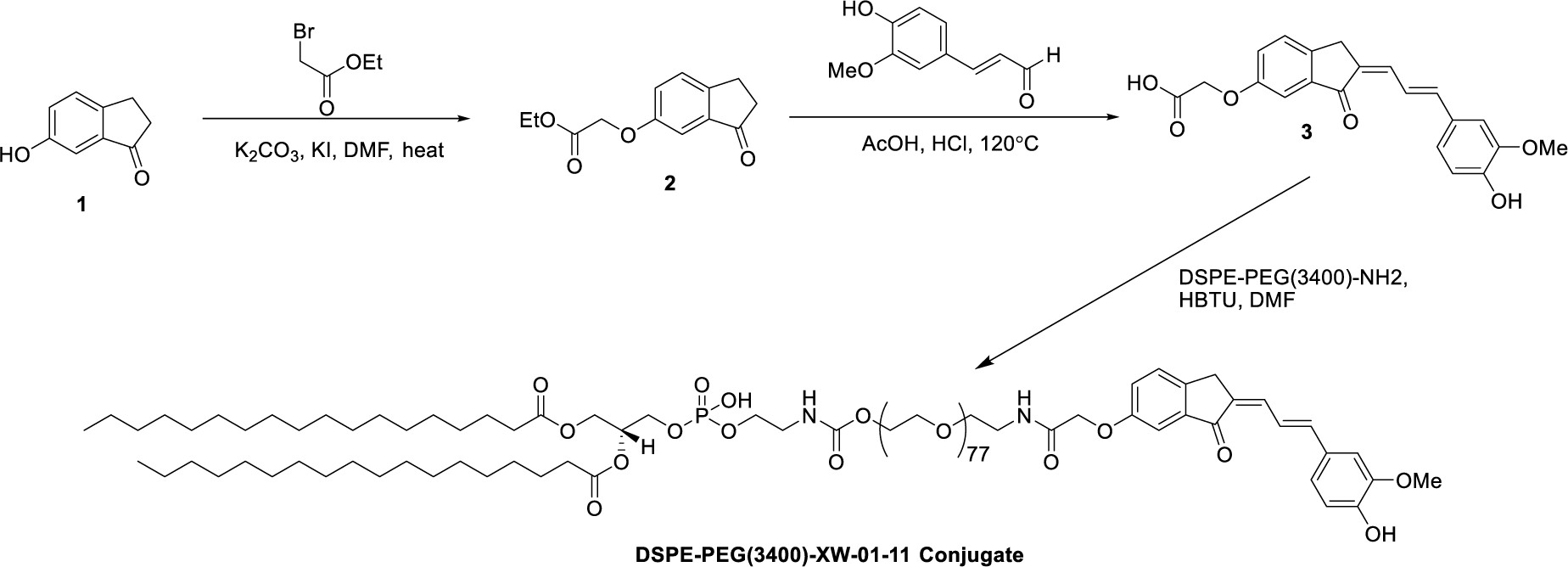
Synthetic route to **DSPE-PEG(3400)-XW-01-11 Conjugate.**

##### Ethyl 2-((3-oxo-2,3-dihydro-1H-inden-5-yl)oxy)acetate (2)

Ethyl bromoacetate (2.3 g, 13.5 mmol) was added in one portion to a solution of 6-hydroxy-1-indanone, **1** (1.0 g, 6.8 mmol), K_2_CO_3_ (2.8 g, 20.2 mmol) and KI (112 mg, 0.7 mmol) in DMF (10 mL). The reaction mixture was stirred at 90 °C overnight. At this point, TLC (silica, 1:3 EtOAc–hexanes) showed that the reaction was complete. The reaction mixture was cooled to room temperature, and then filtered through a pad of celite using ethyl acetate (50 mL) as the eluent and the filtrate concentrated. The residue was purified by column chromatography eluted with ethyl acetate/hexanes gradient to afford the desired product **2** (1.4 g, 90%) as a white solid. ^1^H NMR (600 MHz, CDCl_3_) δ 7.38 (d, *J* = 8.4 Hz, 1H), 7.26 (dd, *J*_1_ = 2.4 Hz, *J*_2_ = 8.4 Hz, 1H), 7.10 (d, *J* = 2.4 Hz, 1H), 4.64 (s, 2H), 4.26 (q, *J* = 7.8 Hz, 2H), 3.06 (t, *J* = 6.0 Hz, 2H), 2.69 (t, *J* = 6.0 Hz, 2H), 1.29 (t, *J* = 7.8 Hz, 3H); ^13^C NMR (150 MHz, CDCl_3_) δ 206.6, 168.3, 157.5, 148.8, 138.1, 127.6, 124.3, 105.7, 65.4, 61.4, 36.8, 25.0, 14.0.

##### 2-{[(Z)-2-((E)-3-(4-Hydroxy-3-methoxyphenyl)allylidene]-3-oxo-2,3-dihydro-1H-inden-5-yl)oxy}acetic acid (3)

A solution of 4-hydroxy-3-methoxy-cinnamaldehyde (588 mg, 3.3 mmol) and compound **2** (700 mg, 3.0 mmol) in acetic acid (10 mL) was slowly added 37% HCl (0.2 mL). The reaction mixture was stirred at 120 °C overnight, and then cooled to room temperature. The cooled solvent was poured into ice water and then filtered out. The solid was recrystallized with methanol to afford the desired product **3** (768 mg, 70%) as a brown solid. ^1^H NMR (600 MHz, DMSO-*d*_6_) δ 13.09 (s, 1H), 9.52 (s, 1H), 7.55 (d, *J* = 8.4 Hz, 1H), 7.32-7.24 (m, 3H), 7.13 (dd, *J*_1_ = 6.0 Hz, *J*_2_ = 9.0 Hz, 1H), 7.08 (d, *J* = 5.4 Hz, 1H), 7.06 (td, *J*_1_ = 3.0 Hz, *J*_2_ = 7.8 Hz, 1H), 6.81 (d, *J* = 8.4 Hz, 1H), 4.79 (s, 2H), 3.85 (s, 3H), 3.84 (s, 2H); ^13^C NMR (150 MHz, DMSO-*d*_6_) δ 192.2, 170.1, 157.6, 148.5, 147.9, 142.9, 142.1, 139.9, 135.3, 133.7, 127.9, 127.5, 123.2, 122.1, 121.9, 115.7, 110.5, 106.3, 64.8, 55.7, 29.5.

##### DSPE-PEG(3400)-XW-01-11 Conjugate

A solution of DSPE-PEG(3400)-NH_2_ (500 mg, 0.1 mmol) and compound **3** (120 mg, 0.3 mmol) in dry DMF (8 mL) was added HSTU (160 mg, 0.4 mmol). The reaction mixture was stirred at room temperature two days overnight. The reaction mixture was concentrated under reduced pressure. The residue was diluted with methanol/water mixture (1:1, 8 ml), and then loaded into a 2000 MWCO dialysis bag and dialyzed against MES buffer (50mM, 2×5 liters) for 8 hours and then water (3×5 liters) for 2 days. The water was then removed by freeze drying to obtain the desired compound as a yellow solid (242 mg, 48%). ^1^H NMR (600 MHz, CDCl_3_) δ 7.45 (d, *J* = 8.4 Hz, 1H), 7.23 (dd, *J*_1_ = 2.4 Hz, *J*_2_ = 5.6 Hz, 1H), 7.19-7.17 (m, 2H), 7.15 (d, *J* = 2.4 Hz, 1H), 7.02–6.99 (m, 3H), 6.78 (dd, *J*_1_ = 8.4 Hz, *J*_2_ = 10.8 Hz, 1H), 5.09 (s, 1H), 4.48 (s, 2H), 4.30 (dd, *J*_1_ = 2.4 Hz, *J*_2_ = 12.0 Hz, 1H), 4.12–4.06 (m, 2H), 4.01 (t, *J* = 4.2 Hz, 2H), 3.98 (s, 2H), 3.81–3.77 (m, 5H), 3.73 (s, 2H), 3.49 (dd, *J*_1_ = 2.4 Hz, *J*_2_ = 7.8 Hz, 2H), 3.48–3.46 (m, 3H), 3.35–3.32 (m, 4H), 3.20–3.18 (m, 2H), 2.21–2.18 (m, 4H), 1.93–1.92 (m, 4H), 1.50–1.47 (m, 4H), 0.761 (t, *J* = 6.0 Hz, 6H); HRMS (MALDI) *calcd* for C_219_H_410_N_2_O_92_P [M+H]^+^ 4571.7203, found, 4571.7069.

##### Fabrication of nano scavengers (T)

Liposomes were prepared using standard hydration/extrusion protocol. Briefly, a lipid mixture containing HSPC, DSPE-mPEG(2000), cholesterol, Gd(III)DSPE-DOTA, DSPE-PEG(3400)-XW-01-11 conjugate prepared at a molar ratio of 32:2.5:40:25:0.5, respectively in ethanol (600 μL) was stirred at 60–65 °C until all of them dissolved to form a clear solution in the ethanol. DHPE-Rhodamine dissolved in ethanol (1 mg in 200 μL) was added to the solution. The dissolved lipids were hydrated with histidine buffered saline (HBS) (10 mM Histidine, 140 mM NaCl, ∼pH 7.6) at 60–65 °C for 45 mins. The hydrated lipid solution was extruded sequentially through 400 nm (5 passes) followed by 200 nm (8 passes) Nuclepore membranes at 60–65 °C using a high-pressure extruder (Northern Lipids, Vancouver, BC, Canada) to form liposomes of desired size. The liposomal suspension was dialyzed against Histidine-buffered saline (HBS) using 300 kDa molecular weight cutoff membranes (Spectrum Laboratories Inc., CA, USA,) to remove any un-encapsulated lipids and ethanol. The control formulation was prepared by the same procedure but the lipid mixture did not include the targeting lipid conjugate, DSPE-PEG(3400)-XW-01-11 conjugate.

##### HMC-3 and T98G cell lines

The human microglia clone 3 cell line (HMC-3) and T98G cells were purchased from American Type Culture Collection (ATCC, Manassas, VA, USA). HMC-3 cells and T98G cells were cultured in Eagle’s Minimum Essential Media (EMEM, ATCC)) supplemented with 10% FBS (Sigma-Aldrich, St. Louis, MO, USA) and 100 units/mL (U/mL) penicillin/streptomycin (Invitrogen, Carlsbad, CA, USA), and incubated in a humidified atmosphere (5% CO_2_) at 37 °C, separately.

##### SHSY5Y Cell

SHSY5Y was purchased from American Tissue Culture Collection (ATCC, Manassas, VA, U.S.A.) and cultured in Dulbecco’s modified Eagle’s medium (DMEM, Thermo Fisher Scientific, Waltham, MA, U.S.A.) with 10% fetal bovine serum (FBS, Thermo Fisher Scientific) and 100 units/mL (U/mL) penicillin/streptomycin (Invitrogen, Carlsbad, CA, USA), and incubated in a humidified atmosphere (5% CO_2_) at 37 °C.

##### Inflammatory potential

(ELISA MAX™ Deluxe Set Human TNF-α; ELISA MAX™ Deluxe Set Human IL-1β; ELISA MAX™ Deluxe Set Human IL-6; Biolegend). Experiments were perfumed described in the technical data sheet.

##### Mice

All animal experiments were conducted under protocols approved by the Institutional Animal Care and Use Committee (IACUC) at Baylor College of Medicine. Animals had free access to food and water.

##### Animal Studies

All animal studies reported in this paper were conducted under study-specific protocols that were specifically approved by the Institutional Animal Care and Use Committee (IACUC) at the Baylor College of Medicine. In all animal experiments, 6 mice were used in each test and control group.

##### Pharmacokinetics and biodistribution study

Images were acquired pre-contrast and then at the following time points post-contrast: 0h (within 5 minutes after injection), 1h, 2h, 4h, 8h, 1 day, 2 days, 4 days, 7 days, 14 days. The formulation was intravenously administered via tail vein at a dose of 0.1 mmol Gd/kg. Images were analyzed in Osirix (version 5.8.5, 64-bit). Region of interests (ROIs) was drawn in blood compartment (inferior vena cava) and select organs (liver, spleen, kidneys, and brain) to generate signal-time curves. Values are reported as T1 relaxation rates (R1) in units of 1/s. Blood half-life (t1/2) was determined by fitting signal change in the IVC to a model of exponential decay and was found to be 20.23 ±0.59 hours.

The PK study was performed in C57BL/6 mice (n=3, 9-12 weeks old). The formulation was intravenously administered via tail vein at a dose of 0.1 mmol Gd (III)/kg. The organs (including liver, spleen, kidney, and brain) were harvested after 14 days, 28 days and 60 days. Gd content of the solution were measured using Inductively Coupled Plasma-Atomic Emission Spectrometry (ICP-AES)

##### Mouse Brain Tissue staining

The mouse brain (perfused by PBS and 4% formalin, stored in 30% sucrose solution) was embedded in Tissue-Tek O.C.T. and kept in the liquid nitrogen for 30 minutes. The embedded tissue was sliced into 30 μm thick sections with Lecia Biosystems Cryostats under −20 °C and mounted onto microscope slides,

##### Anti-alpha-synuclein antibody staining

The brain section was washed with PBS two times and antigen retrieval by citric acid buffer (PH ∼ 6.0)/microwave 30 minutes. Next, the section was washed with PBS three times and permeabilized with 0.1% Triton-X 100 for 10 minutes, followed by a washed with PBS two times. Tissue was incubated with 10% normal goat serum at room temperature for one hour followed by incubation with primary antibody (Anti-Alpha-synuclein (phospho S129) antibody [EP1536Y] (ab51253), Abcam) (1:500 in 1% goat serum) overnight at 4 °C. Tissue was washed with PBS three times and incubated for two hours at room temperature with Alexa Fluor 750 labeled secondary antibody (1:200 in PBS). After washed with PBS two times, tissue was incubated with Hoechst (33342, thermos fisher scientific) at room temperature for 7 minutes. The section was washed with PBS three times, coverslipped, and imaged in Olympus IX81 microscope using standard excitation/emission filters.

##### DAB staining

The brain section was washed with PBS two times and antigen retrieval by citric acid buffer (PH ∼ 6.0)/microwave 30 minutes. The section washed with PBS two times, and then blocked endogenous peroxidase activity by 0.3% H_2_O_2_ for 10 minutes. Next, the section was washed with PBS three times and permeabilized with 0.1% Triton-X 100 for 10 minutes, followed by a washed with PBS three times. The tissue was incubated with 10% normal goat serum at room temperature for one hour followed by incubation with primary antibody (Anti-Alpha-synuclein (phospho S129) antibody [EP1536Y] (ab51253), Abcam) (1:500 in 1% goat serum) overnight at 4 °C. The section was washed with PBS three times and then incubated with a biotinylated secondary antibody (1:200) for 30 minutes, followed by incubation in ABC Elite Kit (1:100 each A and B mixed) for 1 hr. Then, the tissue was washed with PBS three times and incubated in 3,3’-diaminobenzidine solution while checking for staining (1 to 3 minutes approx.), and the reaction was stopped by distill water. The tissue was then dehydrated and cleared with ethanol and xylene, and imaged.

##### Anti-IBA1 antibody staining

The brain section was washed with PBS two times and antigen retrieval by citric acid buffer (PH ∼ 6.0)/microwave 30 minutes. Next, the section was washed with PBS three times and permeabilized with 0.1% Triton-X 100 for 10 minutes, followed by a washed with PBS two times. Tissue was incubated with 10% normal goat serum at room temperature for one hour followed by incubation with primary antibody (IBA1 Polyclonal Antibody, Unconjugated, Host: Rabbit / IgG, Thermo scientific) (1:500 in 1% goat serum) overnight at 4 °C. Tissue was washed with PBS three times and incubated for two hours at room temperature with Alexa Fluor 750 labeled secondary antibody (1:200 in PBS). After washed with PBS two times, tissue was incubated with Hoechst (33342, thermos fisher scientific) at room temperature for 7 minutes. The section was washed with PBS three times, coverslipped, and imaged in Olympus IX81 microscope using standard excitation/emission filters.

##### Anti-GFAP antibody staining

The brain section was washed with PBS two times and antigen retrieval by citric acid buffer (PH ∼ 6.0)/microwave 30 minutes. Next, the section was washed with PBS three times and permeabilized with 0.1% Triton-X 100 for 10 minutes, followed by a washed with PBS two times. Tissue was incubated with 10% normal goat serum at room temperature for one hour followed by incubation with primary antibody (Anti-GFAP Mouse Monoclonal Antibody (clone: 2E1.E9), Biolegend) (1:500 in 1% goat serum) overnight at 4 °C. Tissue was washed with PBS three times and incubated for two hours at room temperature with Alexa Fluor 750 labeled secondary antibody (1:200 in PBS). After washed with PBS two times, tissue was incubated with Hoechst (33342, thermos fisher scientific) at room temperature for 7 minutes. The section was washed with PBS three times, coverslipped, and imaged in Olympus IX81 microscope using standard excitation/emission filters.

##### Anti-Neurofilament H antibody staining

The brain section was washed with PBS two times and antigen retrieval by citric acid buffer (PH ∼ 6.0)/microwave 30 minutes. Next, the section was washed with PBS three times and permeabilized with 0.1% Triton-X 100 for 10 minutes, followed by a washed with PBS two times. Tissue was incubated with 10% normal goat serum at room temperature for one hour followed by incubation with primary antibody (Purified anti-Neurofilament H (NF-H), Phosphorylated [SMI 31], Biolegend) (1:500 in 1% goat serum) overnight at 4 °C. Tissue was washed with PBS three times and incubated for two hours at room temperature with Alexa Fluor 750 labeled secondary antibody (1:200 in PBS). After washed with PBS two times, tissue was incubated with Hoechst (33342, thermos fisher scientific) at room temperature for 7 minutes. The section was washed with PBS three times, coverslipped, and imaged in Olympus IX81 microscope using standard excitation/emission filters.

---

**Figure S1.1.**
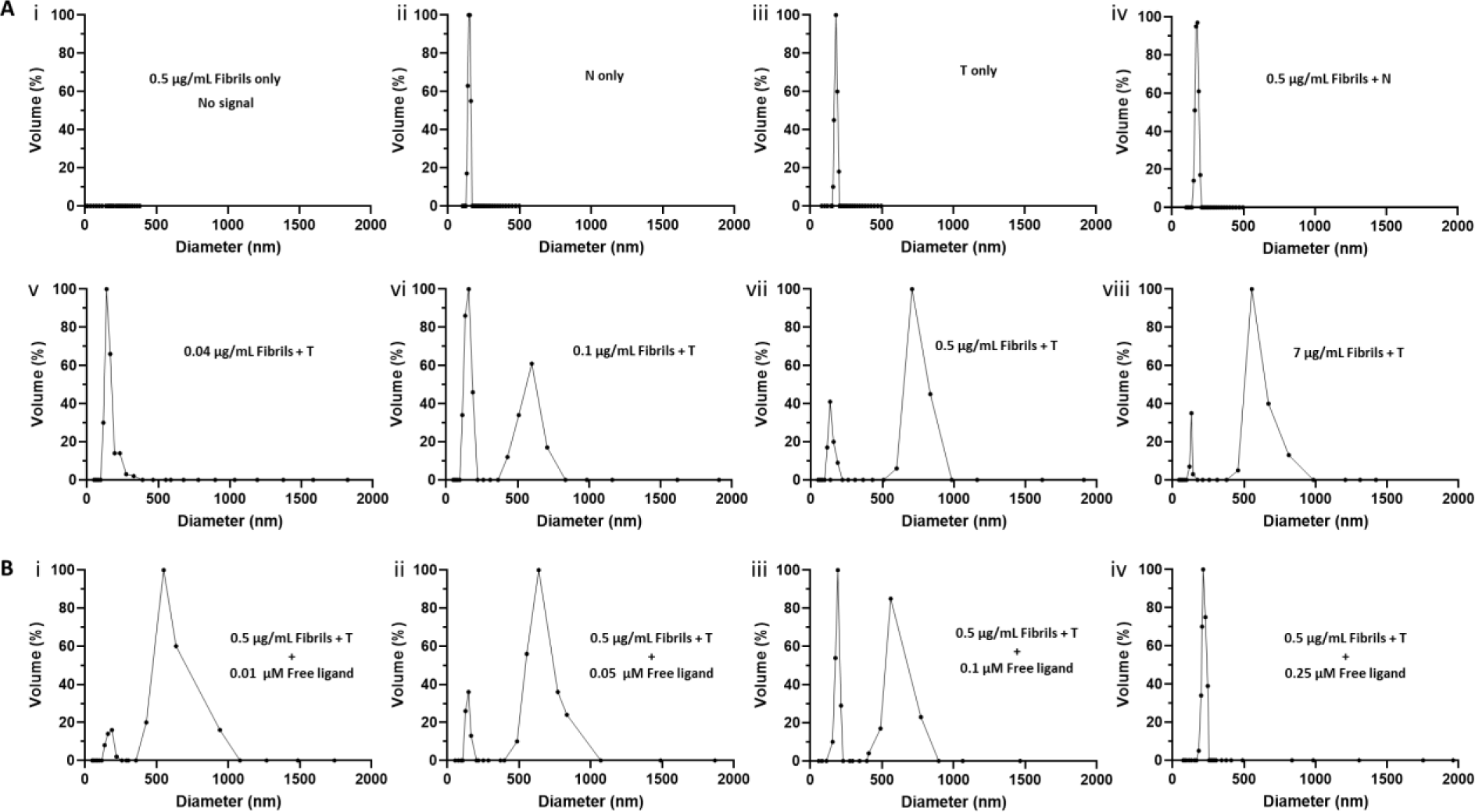
*In vitro* characterization of **T**/α-syn fibril agglomerate formation by DLS. All DLS experiments were run at the same lipid concentration (10 µM lipid) for the **T** and **N** formulations. **A**) DLS profiles of solutions of fibrils at different concentrations with the liposome formulations: **A.i**) A 5 µg/mL solution of synthetic α-fibrils show no DLS signal; **A.ii**) A solution of **N** shows particle distribution of ∼150 nm hydrodynamic diameter; **A.iii**) A solution of **T** shows particle distribution of ∼185 nm hydrodynamic diameter; **A.iv**) A 0.04 µg/mL solution of synthetic α-syn fibrils incubated with **N** shows no major change particle distribution; **A.v**) A 0.04 µg/mL solution of synthetic α-fibrils incubated **T** shows slight distortion at the base of the particle distribution peak; **A.vi**) Increase in fibrils concentration to 0.1 µg/mL results in two different particle populations, the original particle population with a hydrodynamic diameter at ∼185 nm and a new population with a hydrodynamic diameter of ∼600 nm, attributed to **T**/fibril agglomerate formation; **A.vii**) A 0.5 µg/mL solution of synthetic α-fibrils incubated **T** shows two different particle populations, the major particle population with a hydrodynamic diameter of ∼750 nm; **A.viii**) Increase to 7 µg/mL fibrils incubated with **T** shows more decrease in the small particle size distribution. **B)** The blocking experiment were run at the same lipid concentration (10 µM lipid) and α-syn fibrils concentration (0.5 µg/mL), DLS profiles of solutions of free ligand (**XW-01-11**) at different concentrations with the liposome formulations and α-syn fibrils: **B.i**) The free ligand at 0.01 µM concentration shows no major change in the particle distribution; **B.ii**) Increase of the free ligand concentration to 0.05 µM shows an increase in the small particle population suggesting partial blocking of binding site on fibrils; **B.iii**) Further increase of the free ligand concentration to 0.1 µM suggests increase blocking of binding sites on fibrils by the free ligand; **B.iv**) Increase in free ligand concentration to 0.25 µM shows nor large particles form, suggesting complete saturation of binding sites on fibrils.

**Figure S1.2.**
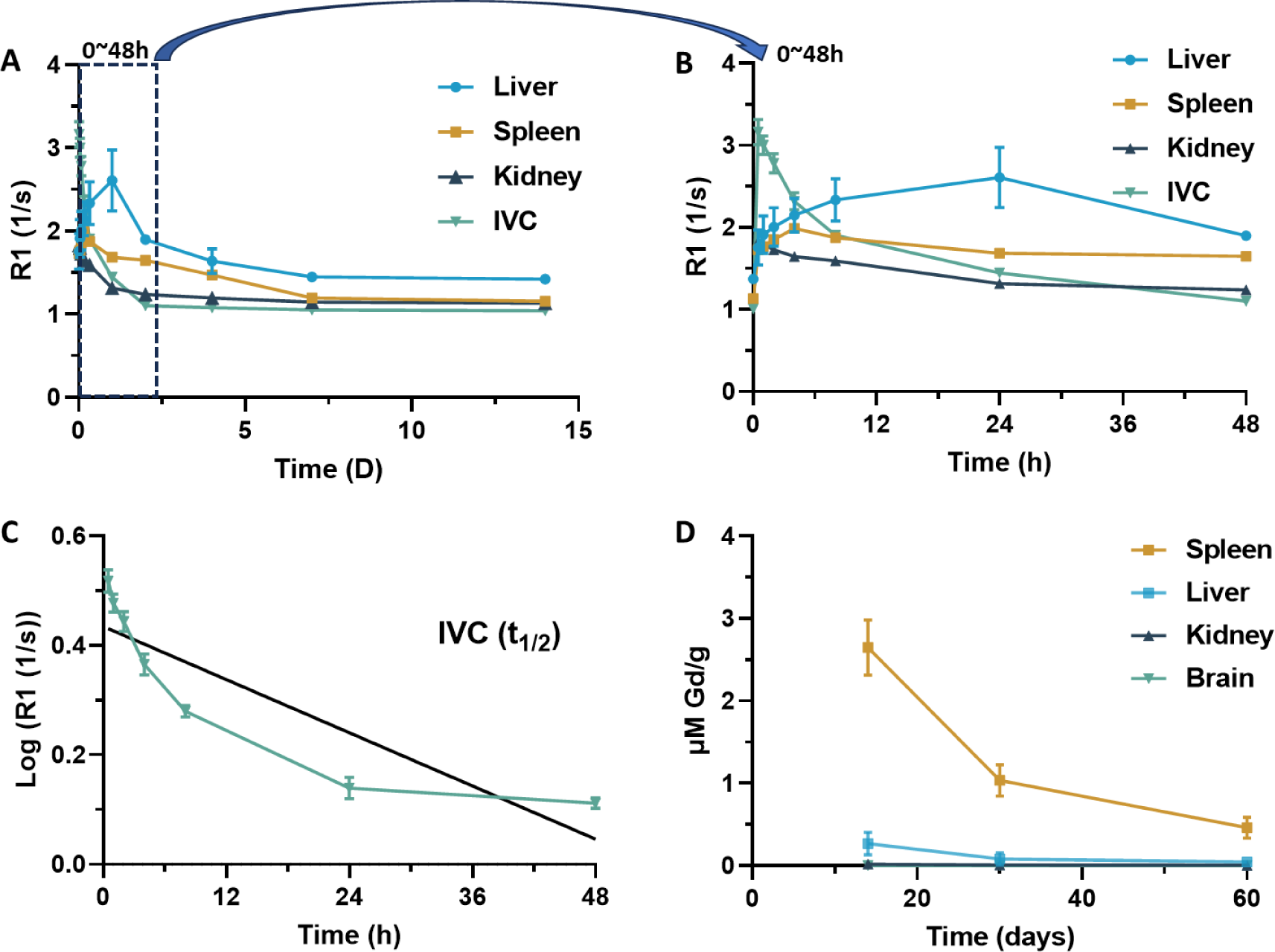
Pharmacokinetics and organ distribution. *In vivo* MRI studies were performed in C57BL/6 mice (n=3, 9-12 weeks old) on a 1T permanent magnet (M2 system, Aspect Imaging, Shoham, Israel) with a 35 mm transmit-receive RF volume coil. **A)** Plot of MRI signal in blood (inferior vena cava) and select organs (liver, spleen, and kidneys) against time over 14 days following contrast injection; **B)** Plot of MRI signal against time between 0 h to 48 hours; **C)** Determination of half-life (t_1/2_); **D)** Distribution of Gd(III) in select organs (liver, spleen, brain, kidney) from 14 days to 60 days determined by ICP analysis of tissue.

**Figure S1.3.**
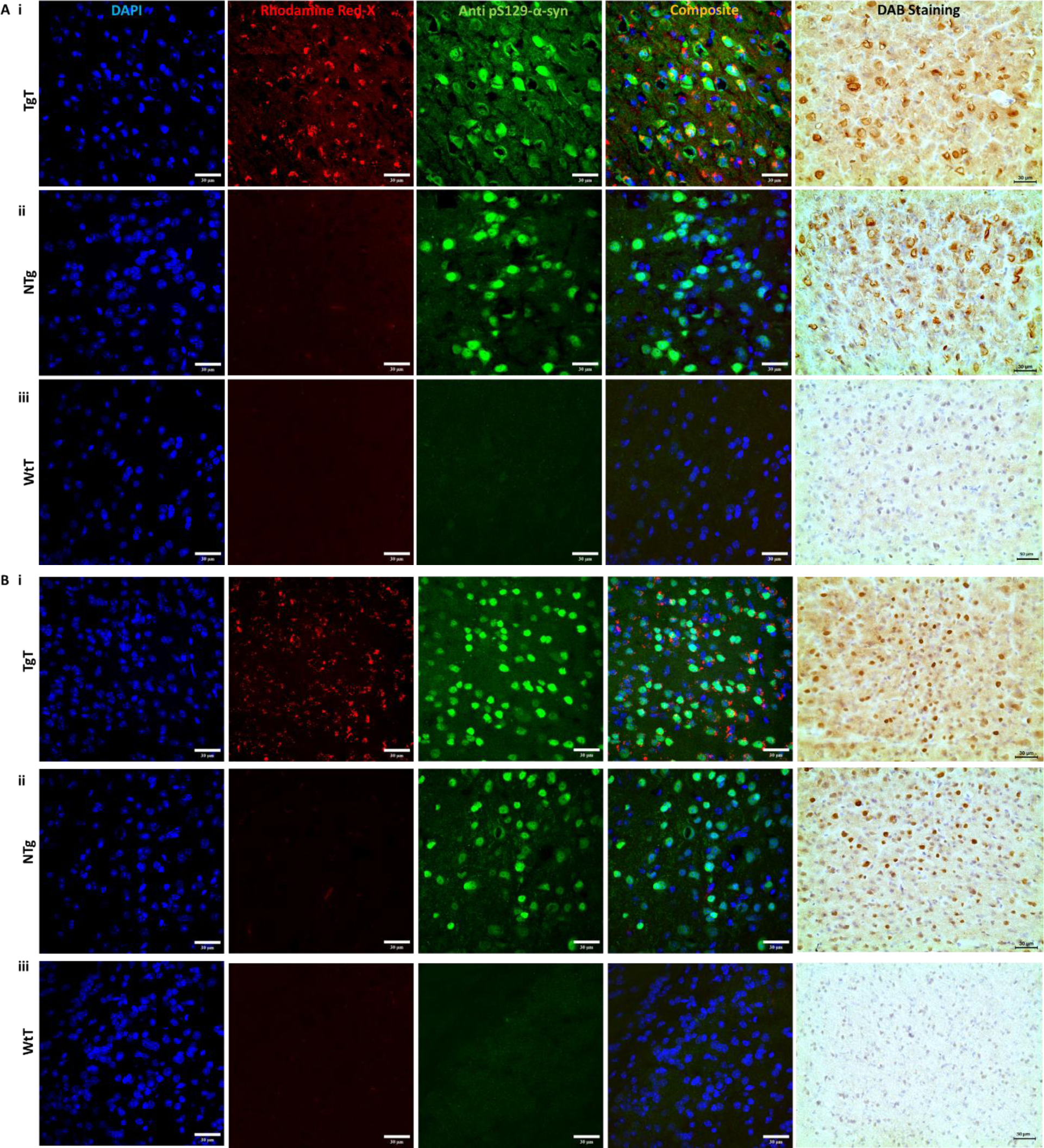
Immunohistochemical analysis demonstrates a correlation between *in vivo* MRI signal enhancement, nano scavenger location and Lewy pathology distribution. **A**) Anti-pSer129-α-syn stained **olfactory bulb** tissue sections from a 16-19 month-old mouse: **A.i) TgT** mouse shows colocalization of **T** (Rhodamine fluorescence) with anti pS129-α-syn antibody signal (Lewy pathology). Lewy pathology is confirmed with DAB staining; **A.ii) TgN** mouse shows little to no Rhodamine fluorescence signal (consistent with *in vivo* MRI), but strong pS129-α-syn pathologic structures (transgenic mouse); **A.iii) WtT** mouse shows no Rhodamine fluorescence signal and no anti pS129-α-syn activity; **B**) Anti-pSer129-α-syn stained olfactory bulb tissue sections from a 13-15-month-old mice similar patterns of Lewy pathology patterns, albeit less dense. Scale bar, 30 μm.

**Figure S1.4.**
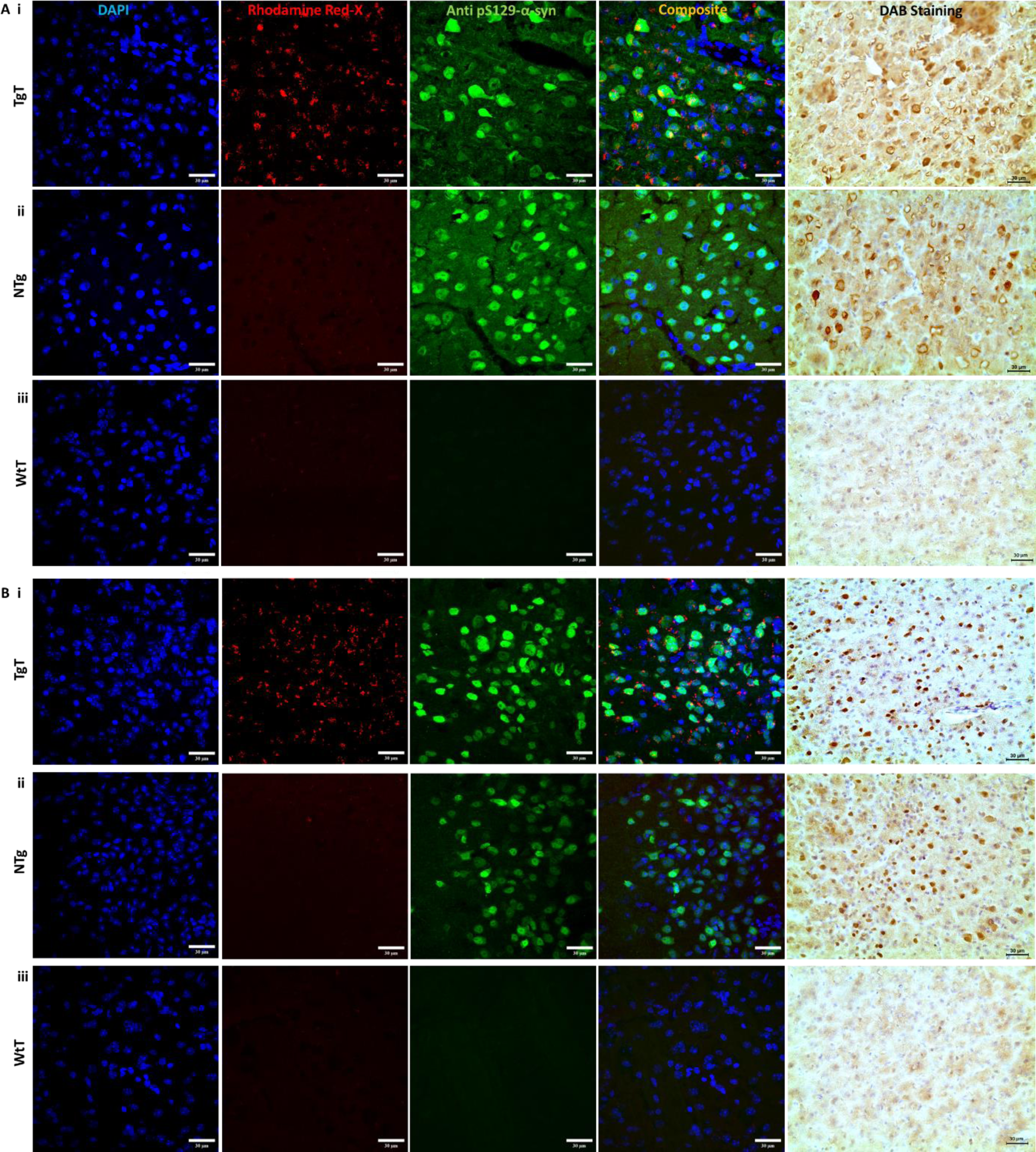
Immunohistochemical analysis demonstrates a correlation between *in vivo* MRI signal enhancement, nano scavenger location and Lewy pathology distribution. **A**) Anti-pSer129-α-syn stained **entorhinal cortex** tissue sections from a 16-19 month-old mouse: **A.i) TgT** mouse shows colocalization of **T** (Rhodamine fluorescence) with anti pS129-α-syn antibody signal (Lewy pathology). Lewy pathology is confirmed with DAB staining; **A.ii) TgN** mouse shows little to no Rhodamine fluorescence signal (consistent with *in vivo* MRI), but strong pS129-α-syn pathologic structures (transgenic mouse); **A.iii) WtT** mouse shows no Rhodamine fluorescence signal and no anti pS129-α-syn activity; **B**) Anti-pSer129-α-syn stained entorhinal cortex tissue sections from a 13-15-month-old mice similar patterns of Lewy pathology patterns, albeit less dense. Scale bar, 30 μm.

**Figure S1.5.**
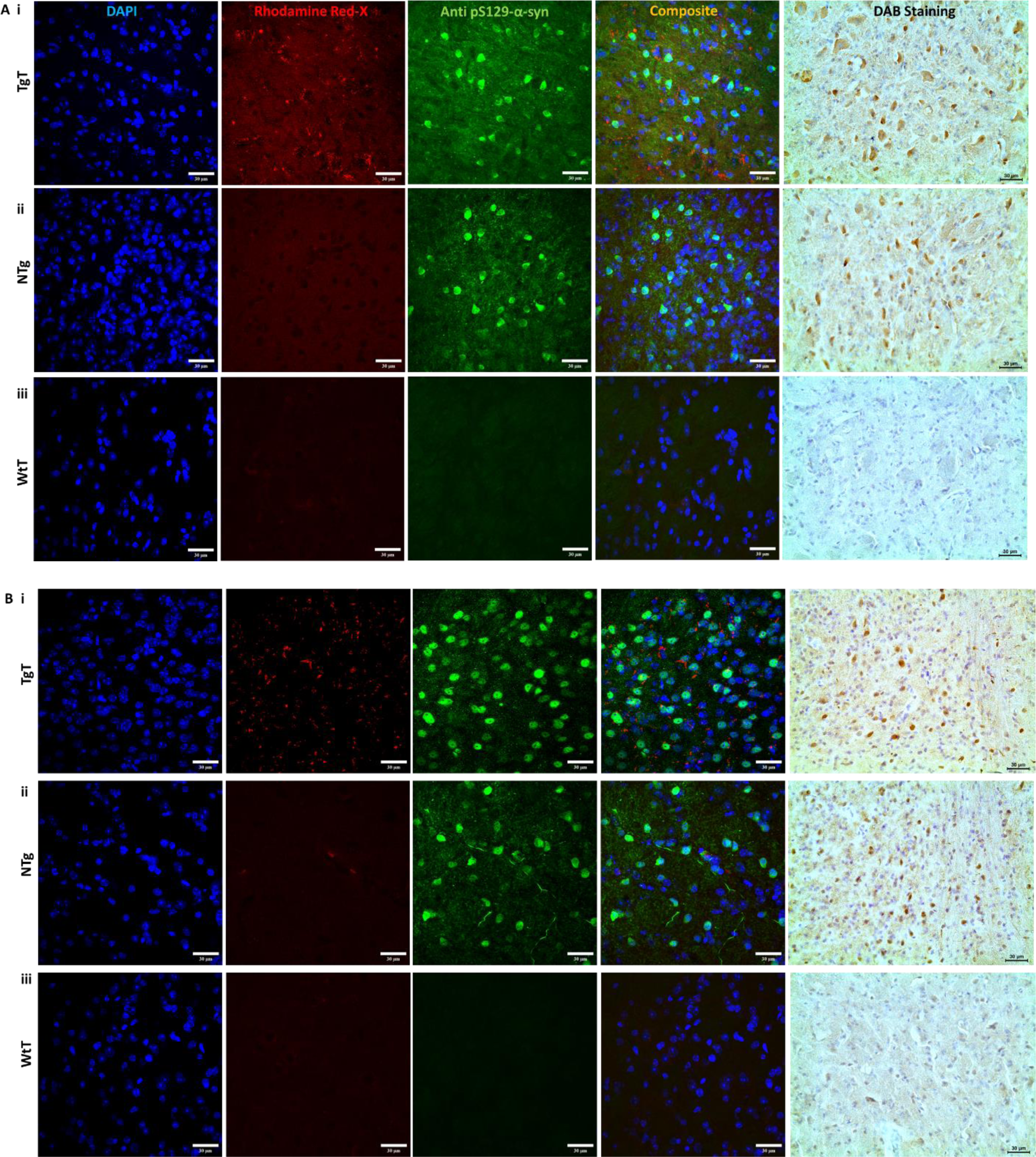
Immunohistochemical analysis demonstrates a correlation between *in vivo* MRI signal enhancement, nano scavenger location and Lewy pathology distribution. **A**) Anti-pSer129-α-syn stained **brainstem** tissue sections from a 16-19 month-old mouse: **A.i) TgT** mouse shows colocalization of **T** (Rhodamine fluorescence, red) with anti pS129-α-syn antibody signal (Lewy pathology). Lewy pathology is confirmed with DAB staining; **A.ii) TgN** mouse shows little to no Rhodamine fluorescence signal (consistent with *in vivo* MRI), but strong pS129-α-syn pathologic structures (transgenic mouse); **A.iii) WtT** mouse shows no Rhodamine fluorescence signal and no anti pS129-α-syn activity; **B**) Anti-pSer129-α-syn stained brainstem tissue sections from a 13-15-month-old mice similar patterns of Lewy pathology patterns, albeit less dense. Scale bar, 30 μm.

**Figure S1.6.**
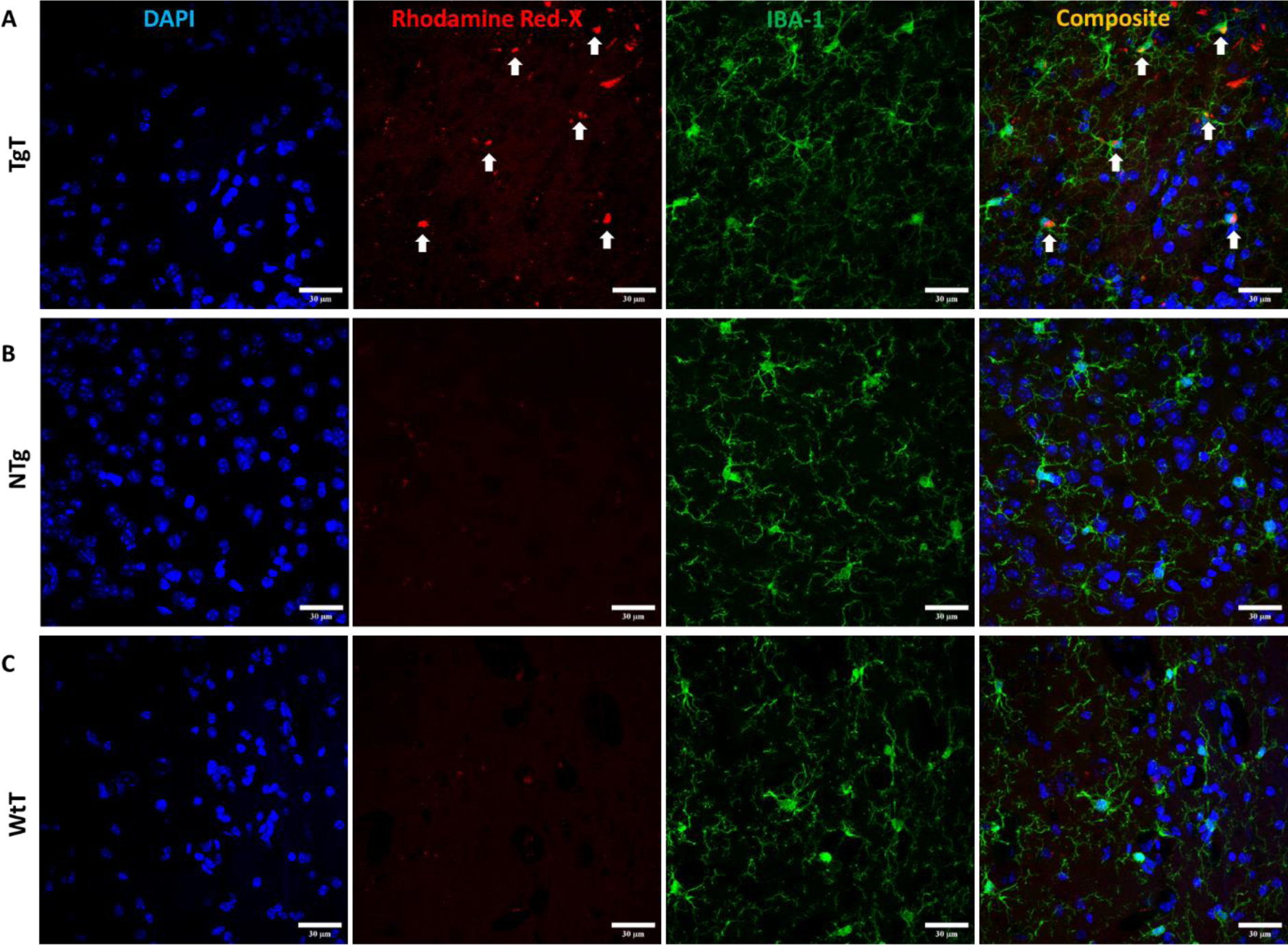
Microglia display *in vivo* uptake of nano scavenger species with apparent compact structural formation (white arrows) in the **Olfactory bulb** as 16–19-month-old **TgT**, **TgN** and **WtT** mice all show cell bodies and processes with strong IBA-1 reactivity **A)** Composite images from imaged **TgT** tissue sections colocalization of NS signal (Rhodamine) in cell bodies of anti IBA-1 reactive cells; Controls **B** and **C** (**TgN** and **WtT** respectively) sections show very faint to no signal in the Rhodamine channel. Scale bar, 30 μm.

**Figure S1.7.**
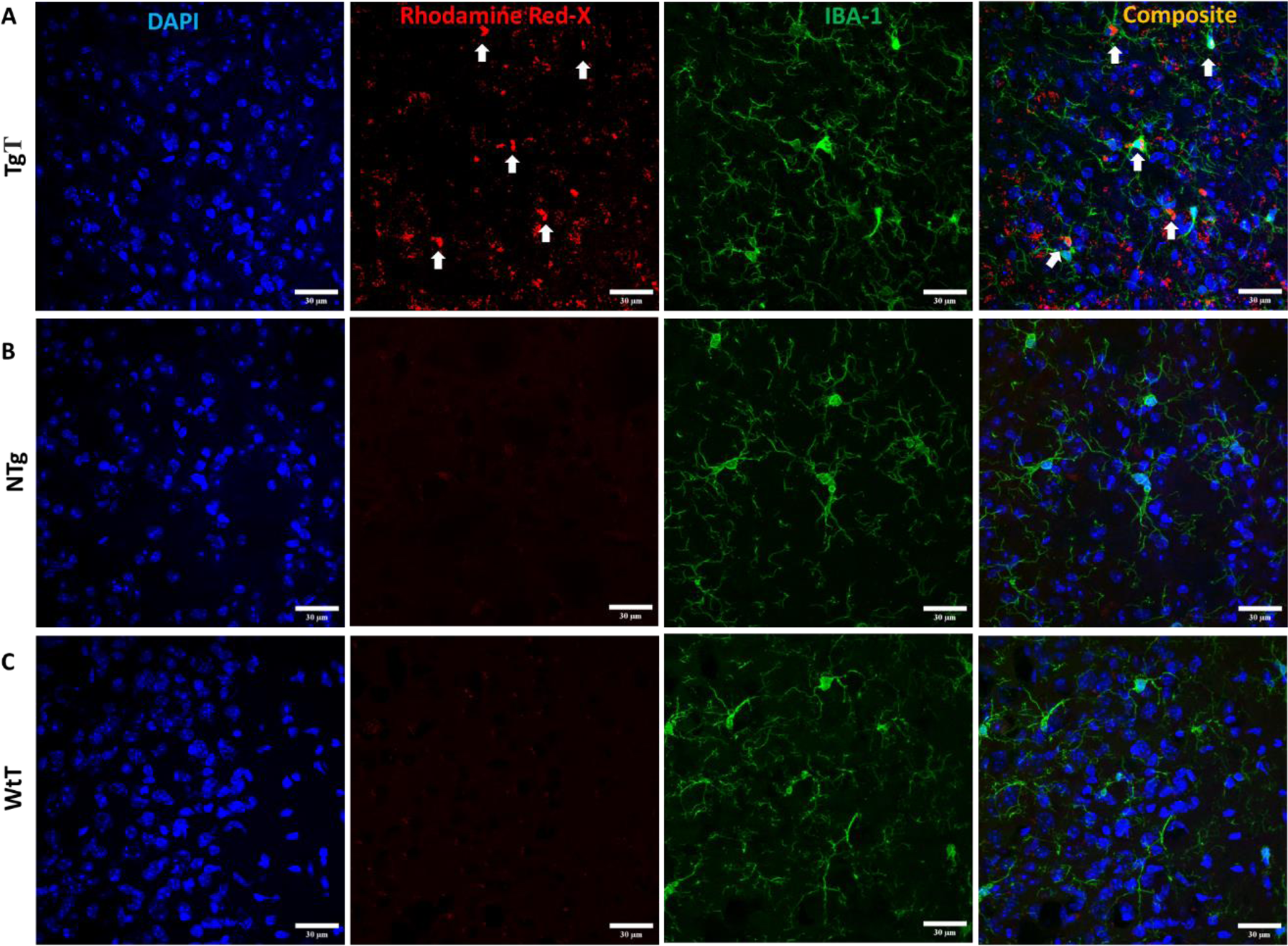
Microglia display *in vivo* uptake of nano scavenger species with apparent compact structural formation (white arrows) in the **Entorhinal cortex** as 16–19-month-old **TgT**, **TgN** and **WtT** mice all show cell bodies and processes with strong IBA-1 reactivity **A)** Composite images from imaged **TgT** tissue sections colocalization of NS signal (Rhodamine) in cell bodies of anti IBA-1 reactive cells; Controls **B** and **C** (**TgN** and **WtT** respectively) sections show very faint to no signal in the Rhodamine channel. Scale bar, 30 μm.

**Figure S1.8.**
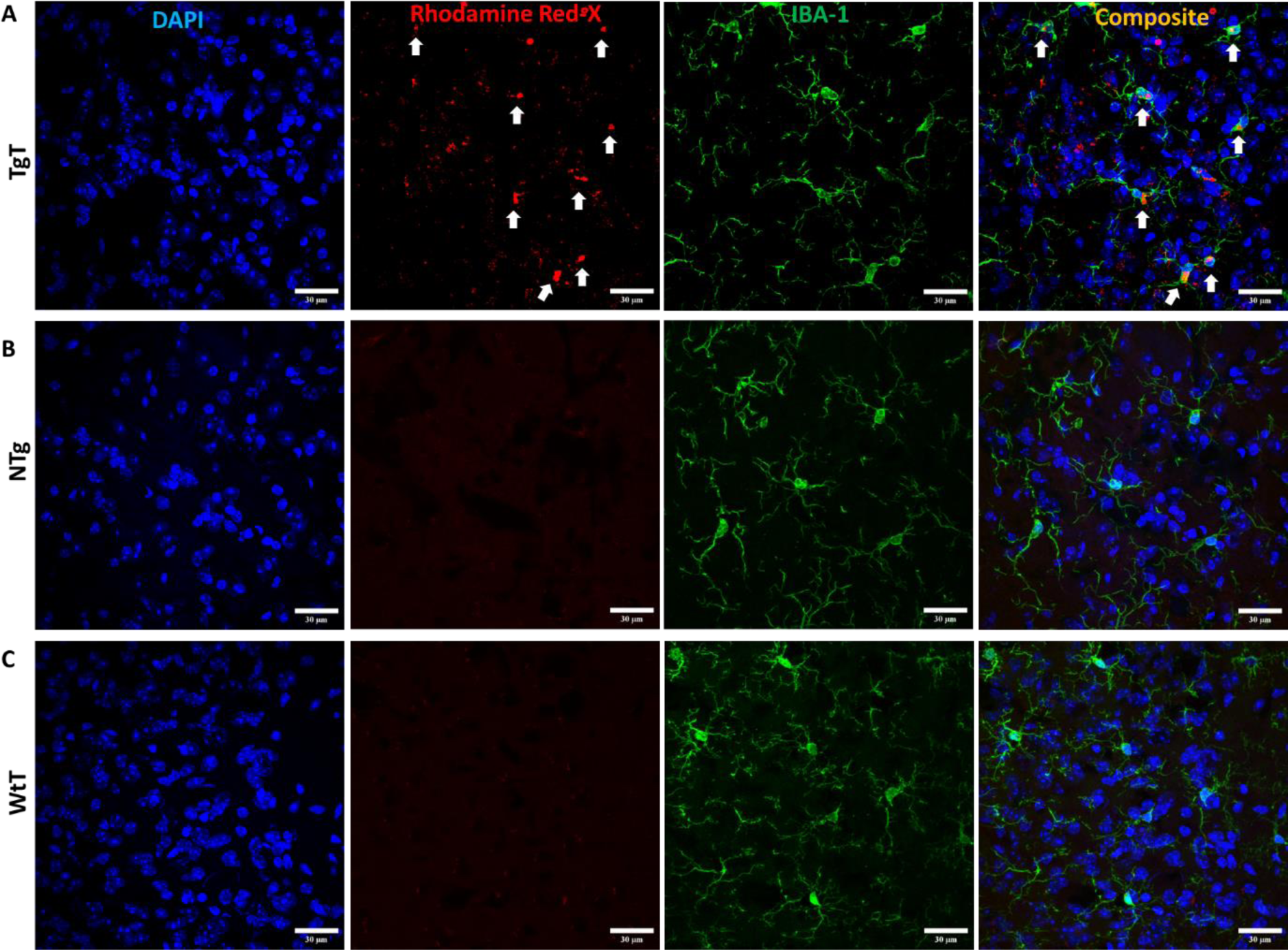
Microglia display *in vivo* uptake of nano scavenger species with apparent compact structural formation (white arrows) in the **Brainstem** as 16–19-month-old **TgT**, **TgN** and **WtT** mice all show cell bodies and processes with strong IBA-1 reactivity **A)** Composite images from imaged **TgT** tissue sections colocalization of NS signal (Rhodamine) in cell bodies of anti IBA-1 reactive cells; Controls **B** and **C** (**TgN** and **WtT** respectively) sections show very faint to no signal in the Rhodamine channel. Scale bar, 30 μm.

**Figure S1.9..**
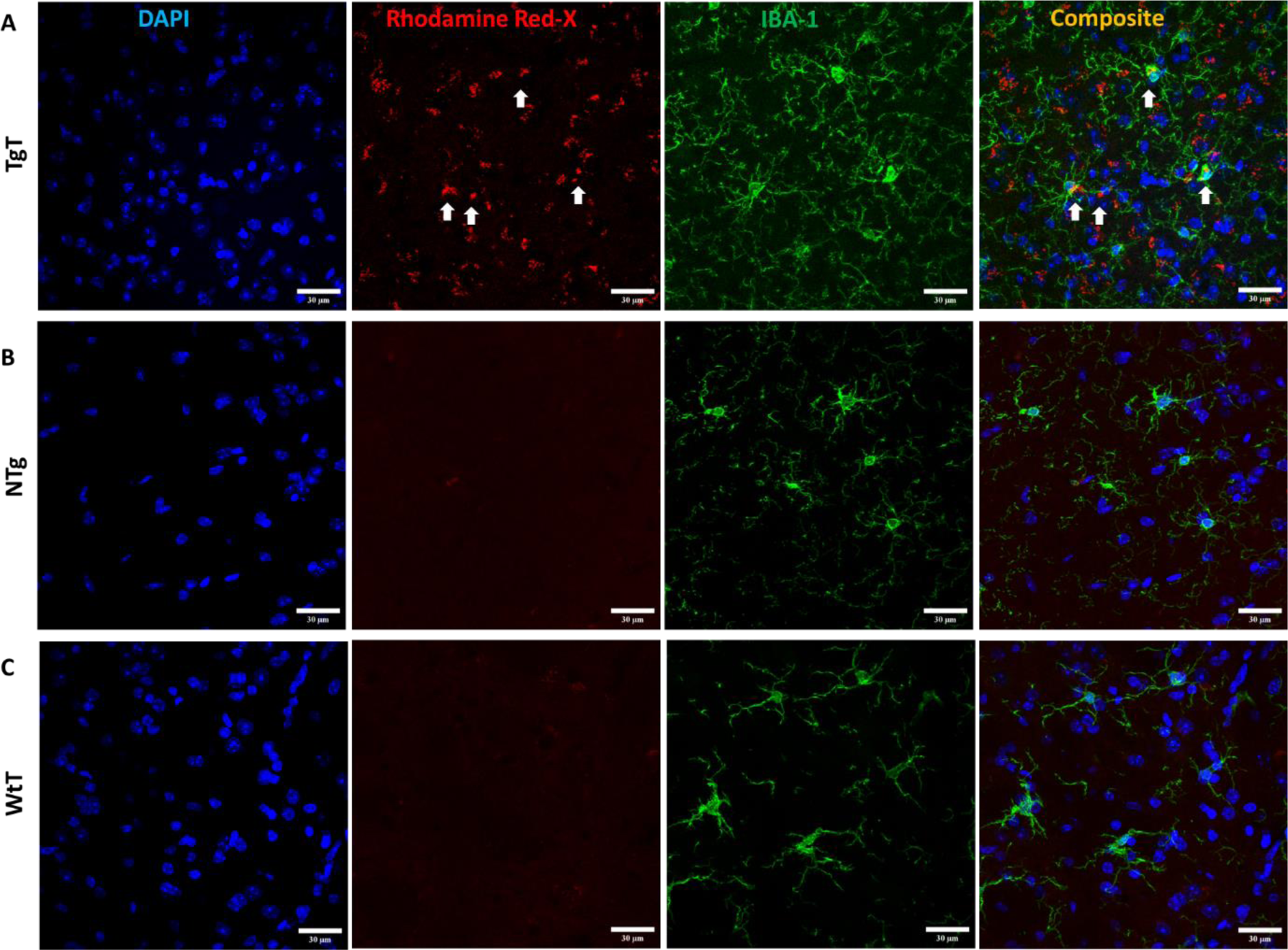
Microglia display *in vivo* uptake of nano scavenger species with apparent compact structural formation (white arrows) in the **Olfactory bulb** as 13–15-month-old **TgT**, **TgN** and **WtT** mice all show cell bodies and processes with strong IBA-1 reactivity **A)** Composite images from imaged **TgT** tissue sections colocalization of NS signal (Rhodamine) in cell bodies of anti IBA-1 reactive cells; Controls **B** and **C** (**TgN** and **WtT** respectively) sections show very faint to no signal in the Rhodamine channel. Scale bar, 30 μm.

**Figure S1.10..**
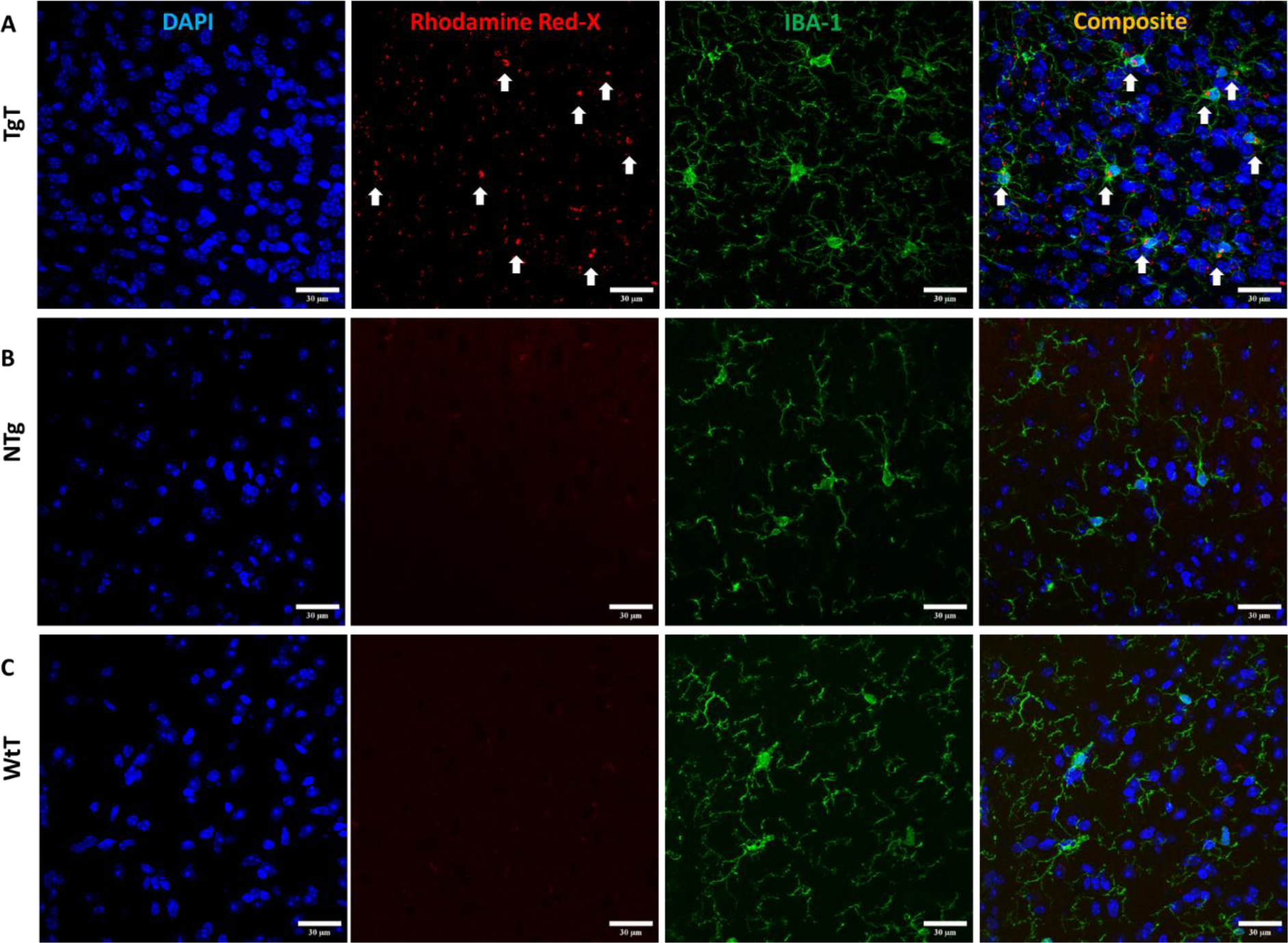
Microglia display *in vivo* uptake of nano scavenger species with apparent compact structural formation (white arrows) in the **Entorhinal cortex** as 13–15-month-old **TgT**, **TgN** and **WtT** mice all show cell bodies and processes with strong IBA-1 reactivity **A)** Composite images from imaged **TgT** tissue sections colocalization of NS signal (Rhodamine) in cell bodies of anti IBA-1 reactive cells; Controls **B** and **C** (**TgN** and **WtT** respectively) sections show very faint to no signal in the Rhodamine channel. Scale bar, 30 μm.

**Figure S1.11..**
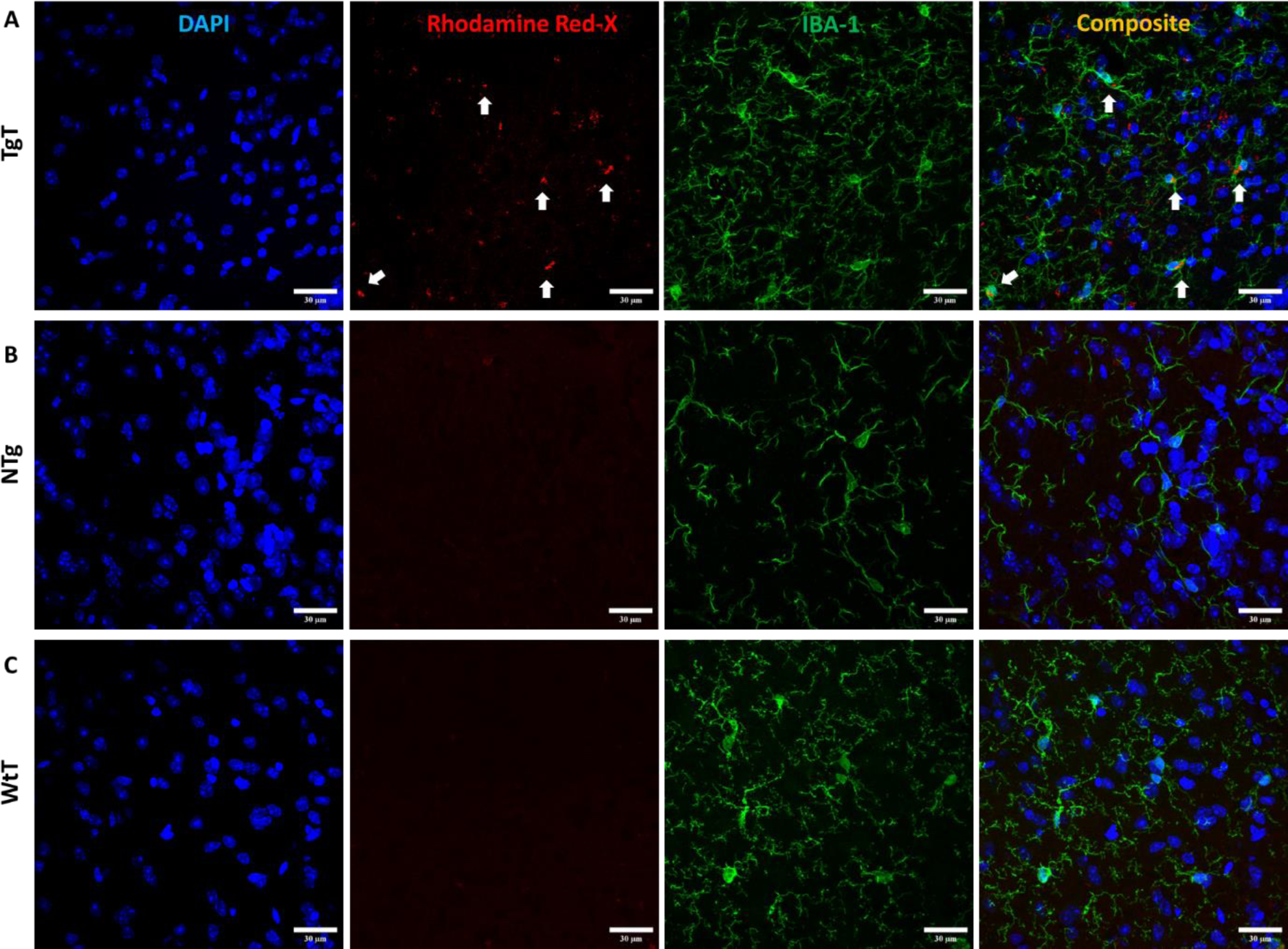
Microglia display *in vivo* uptake of nano scavenger species with apparent compact structural formation (white arrows) in the **Brainstem** as 13–15-month-old **TgT**, **TgN** and **WtT** mice all show cell bodies and processes with strong IBA-1 reactivity **A)** Composite images from imaged **TgT** tissue sections colocalization of NS signal (Rhodamine) in cell bodies of anti IBA-1 reactive cells; Controls **B** and **C** (**TgN** and **WtT** respectively) sections show very faint to no signal in Rhodamine channel. Scale bar, 30 μm.

**Figure S1.12.**
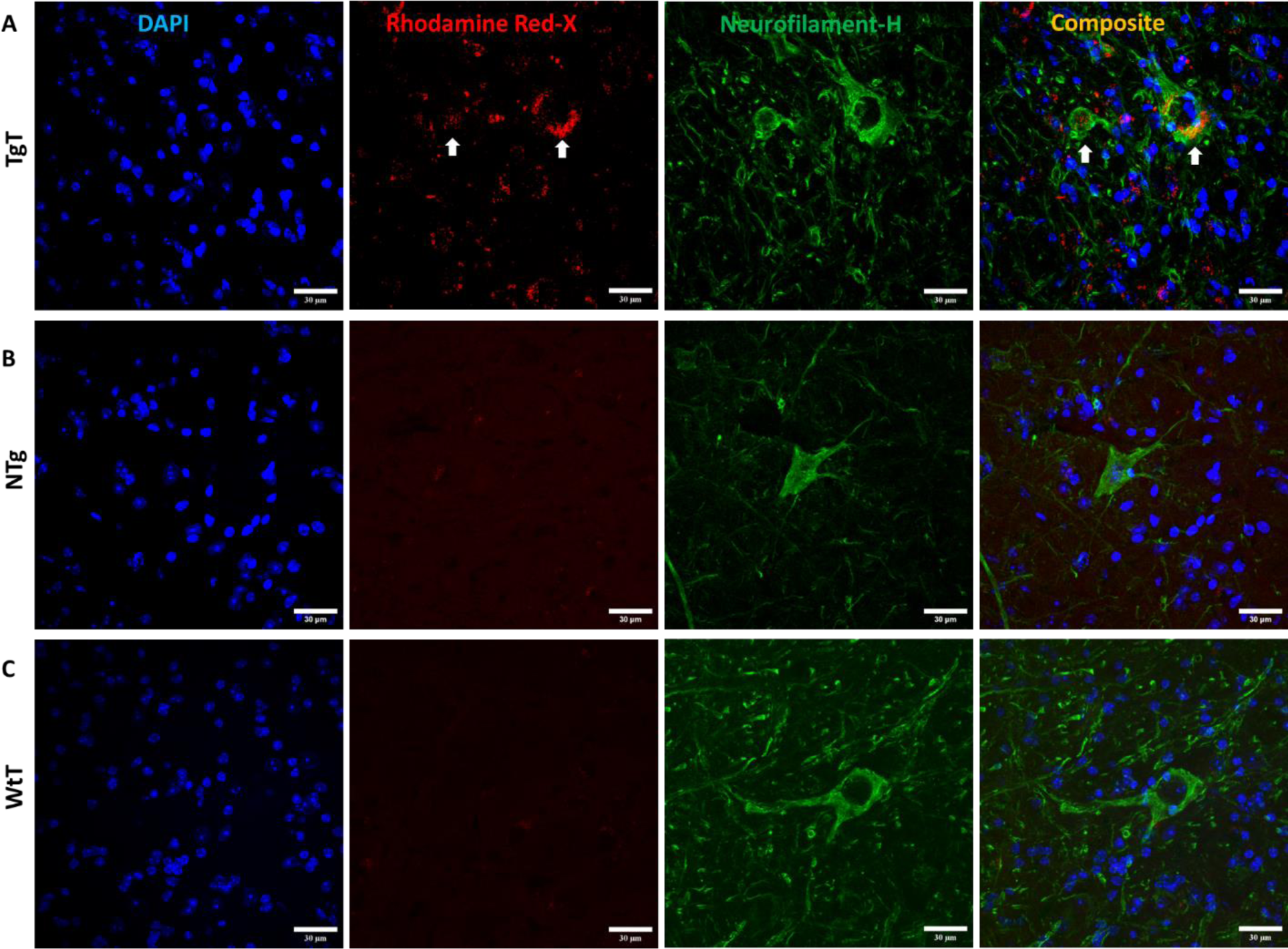
Neuronal cell bodies *in vivo* uptake of nano scavenger species with apparent diffuse structural formation (white arrows). **Brainstem** sections from 16-19-month-old **TgT**, **TgN** and **WtT** mice show cell bodies and processes with strong anti-NF-H reactivity (green fluorescence). **A)** Composite images of **TgT** sections show that the diffuse Rhodamine signal is cytosolic (white arrows). **B)** and **C) TgN** and **WtT** sections respectively show no signal in the Rhodamine channel. Scale bar, 30 μm.

**Figure S1.13.**
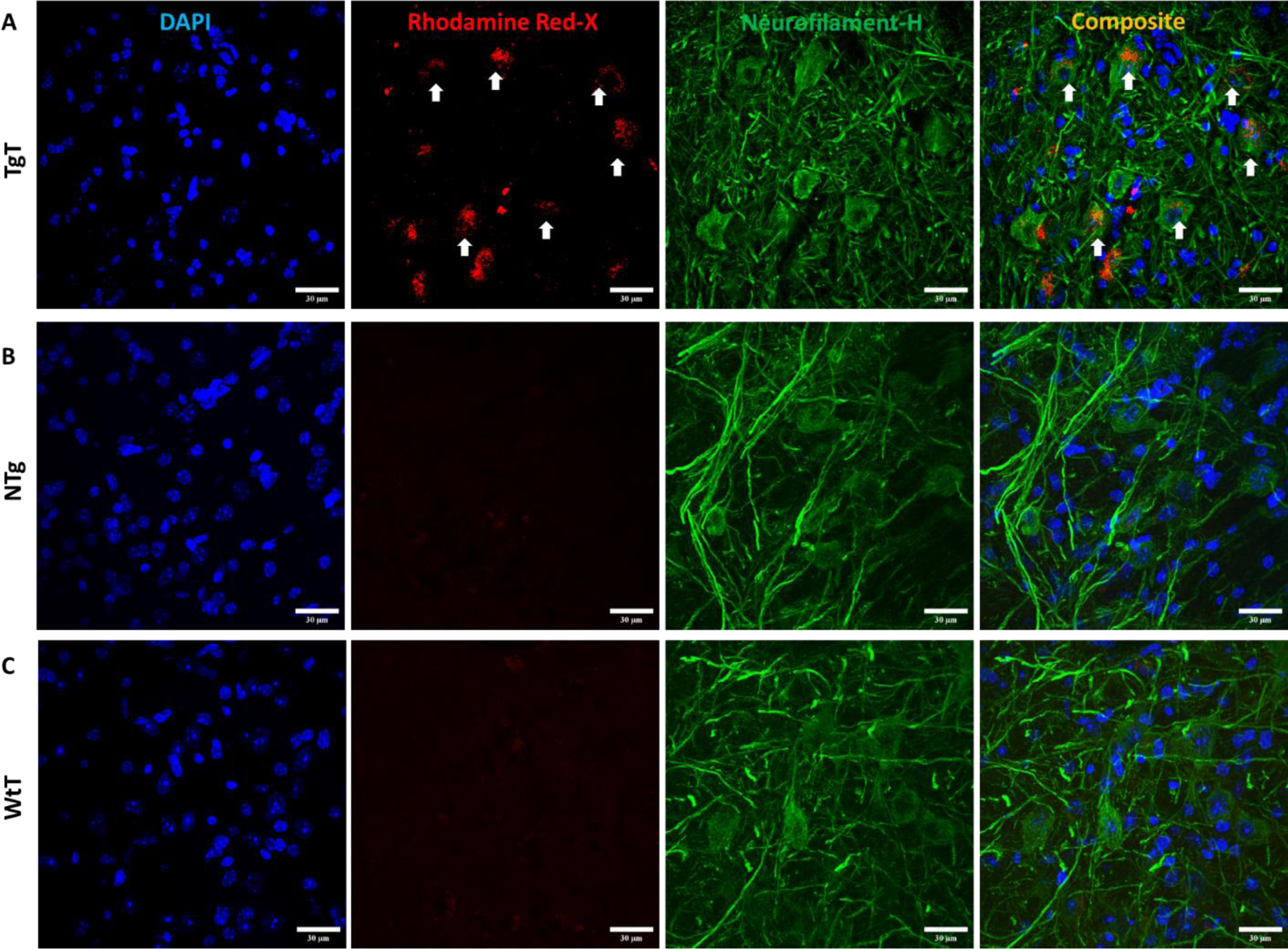
Neuronal cell bodies *in vivo* uptake of nano scavenger species with apparent diffuse structural formation. **Brainstem** sections from 13-15-month-old **TgT**, **TgN** and **WtT** mice show cell bodies and processes with strong anti-NF-H reactivity (green fluorescence). **A)** Composite images shows that the diffuse Rhodamine signal is cytosolic (white arrows). **B**) **TgN** and **C**) **WtT** sections show low to no signal in Rhodamine channel. Scale bar, 30 μm.

**Figure S1.14.**
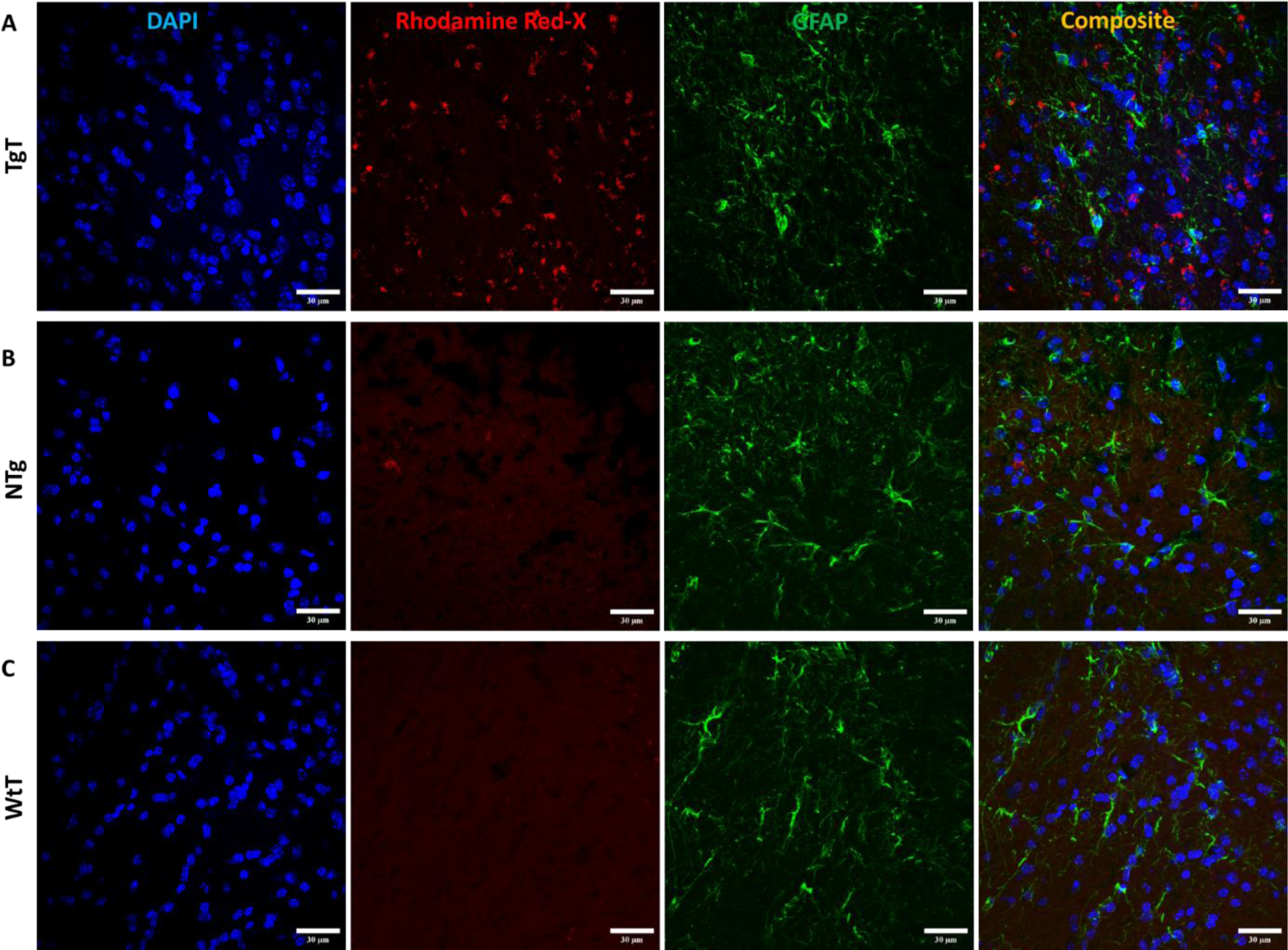
Astrocytes do not display convincing evidence of *in vivo* uptake. **Olfactory bulb** sections from 16-19-month-old **TgT**, **TgN** and **WtT** mice show cell bodies and processes with strong anti GFAP reactivity (green fluorescence). **A) TgT** sections show strong NS signal (Rhodamine), but composite images do not show conclusive evidence of internalization of **T** by GFAP reactive cells. **B) TGN** and **C)WtT** sections show no signal in the Rhodamine channel. Scale bar, 30 μm.

**Figure S1.15.**
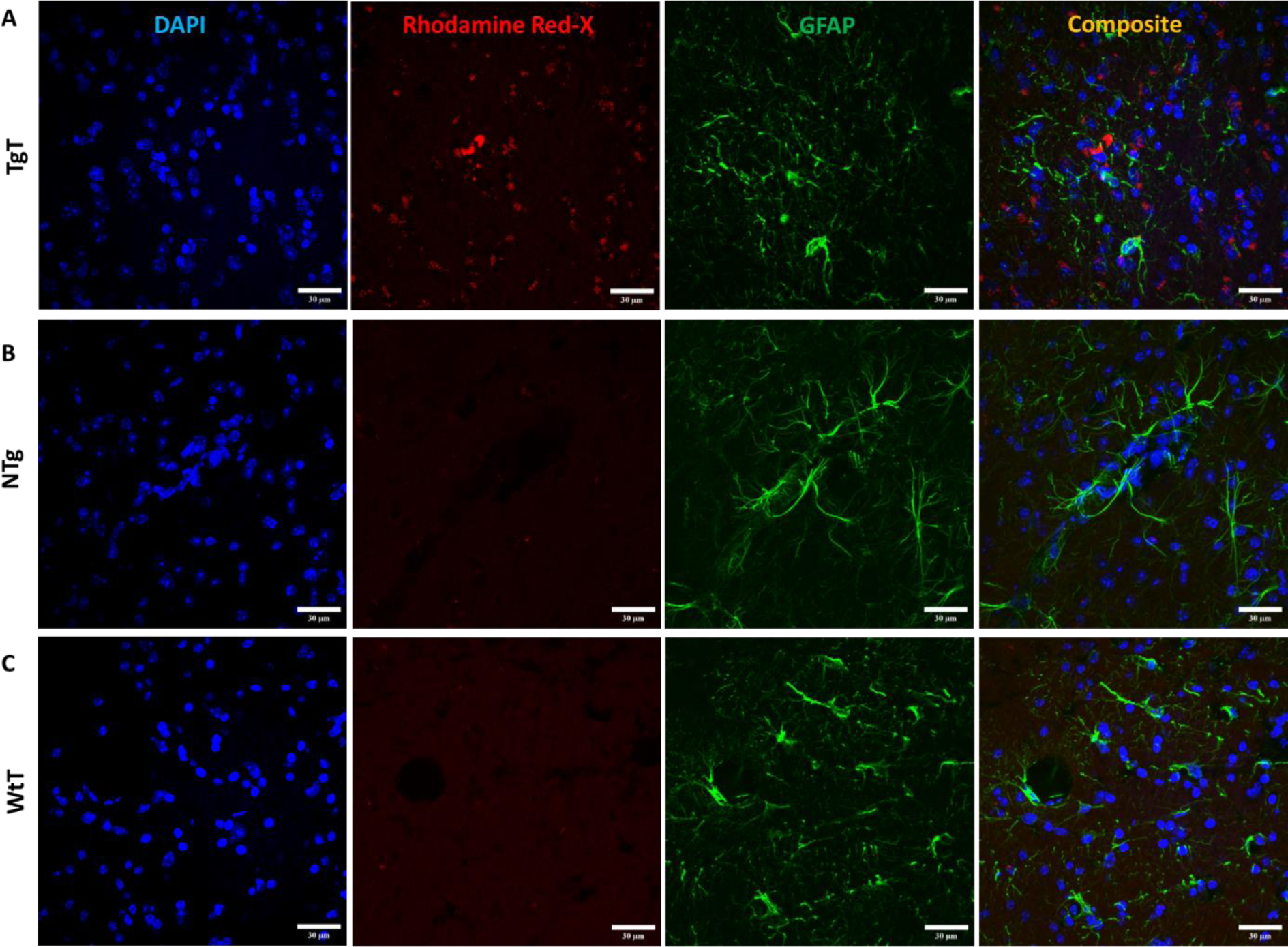
Astrocytes do not display convincing evidence of *in vivo* uptake. Cortex sections from 16-19-month-old **TgT**, **TgN** and **WtT** mice show cell bodies and processes with strong GFAP reactivity (green fluorescence). **A) TgT** sections show strong NS signal (Rhodamine), but composite images do not show conclusive evidence of internalization of **T** by anti GFAP reactive cells. **B) TgN** and **C) WtT** sections show no signal in the Rhodamine channel. Scale bar, 30 μm.

**Figure S1.16.**
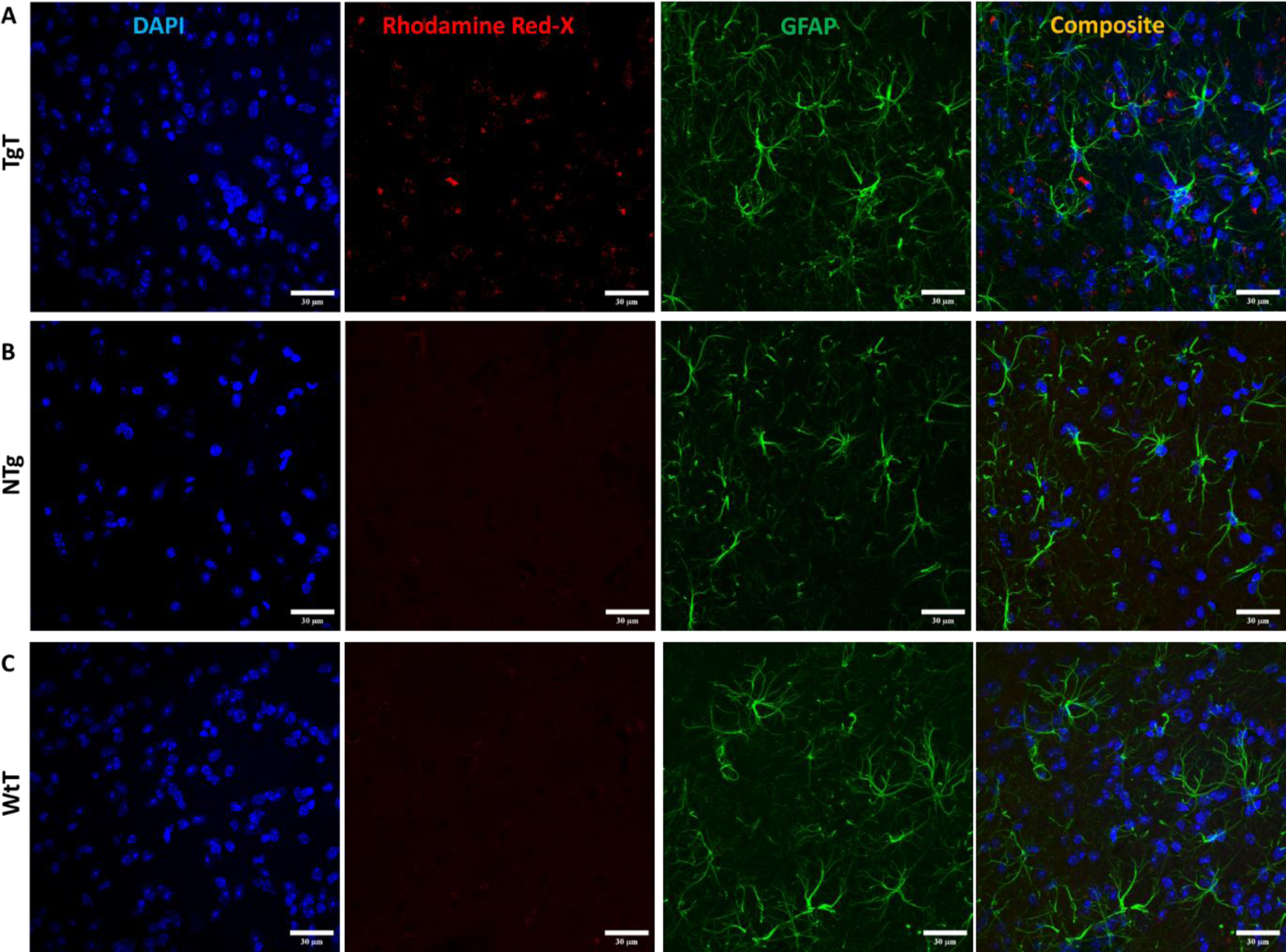
Astrocytes do not display convincing evidence of *in vivo* uptake. **Brainstem** sections from 16-19-month-old **TgT**, **TgN** and **WtT** mice show cell bodies and processes with strong GFAP reactivity (green fluorescence). **A) TgT** sections show strong NS signal (Rhodamine), but composite images do not show conclusive evidence of internalization of **T** by GFAP reactive cells. **B) TgN** and **C) WtT** sections show very low to no signal in Rhodamine channel. Scale bar, 30 μm.

**Figure S1.17.**
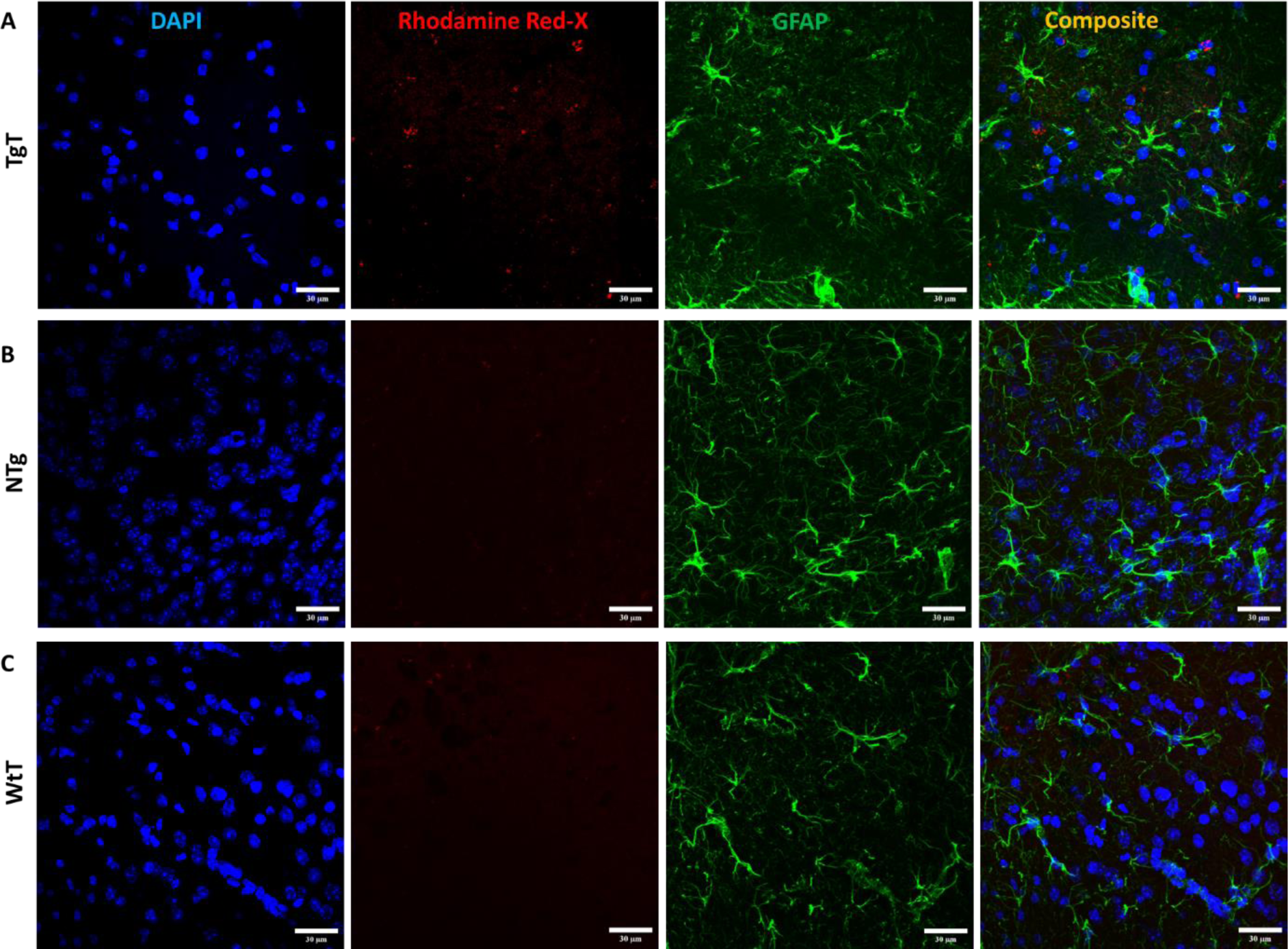
Astrocytes do not display convincing evidence of *in vivo* uptake. **Olfactory bulb** sections from 13-15-month-old **TgT**, **TgN** and **WtT** mice show cell bodies and processes with strong GFAP reactivity. **A) TgT** sections show strong NS signal (Rhodamine) but composite images do not show conclusive evidence of internalization of **T** by GFAP reactive cells. **B) TgN** and **C**) **WtT** sections show very low to no signal in the Rhodamine channel. Scale bar, 30 μm.

**Figure S1.18.**
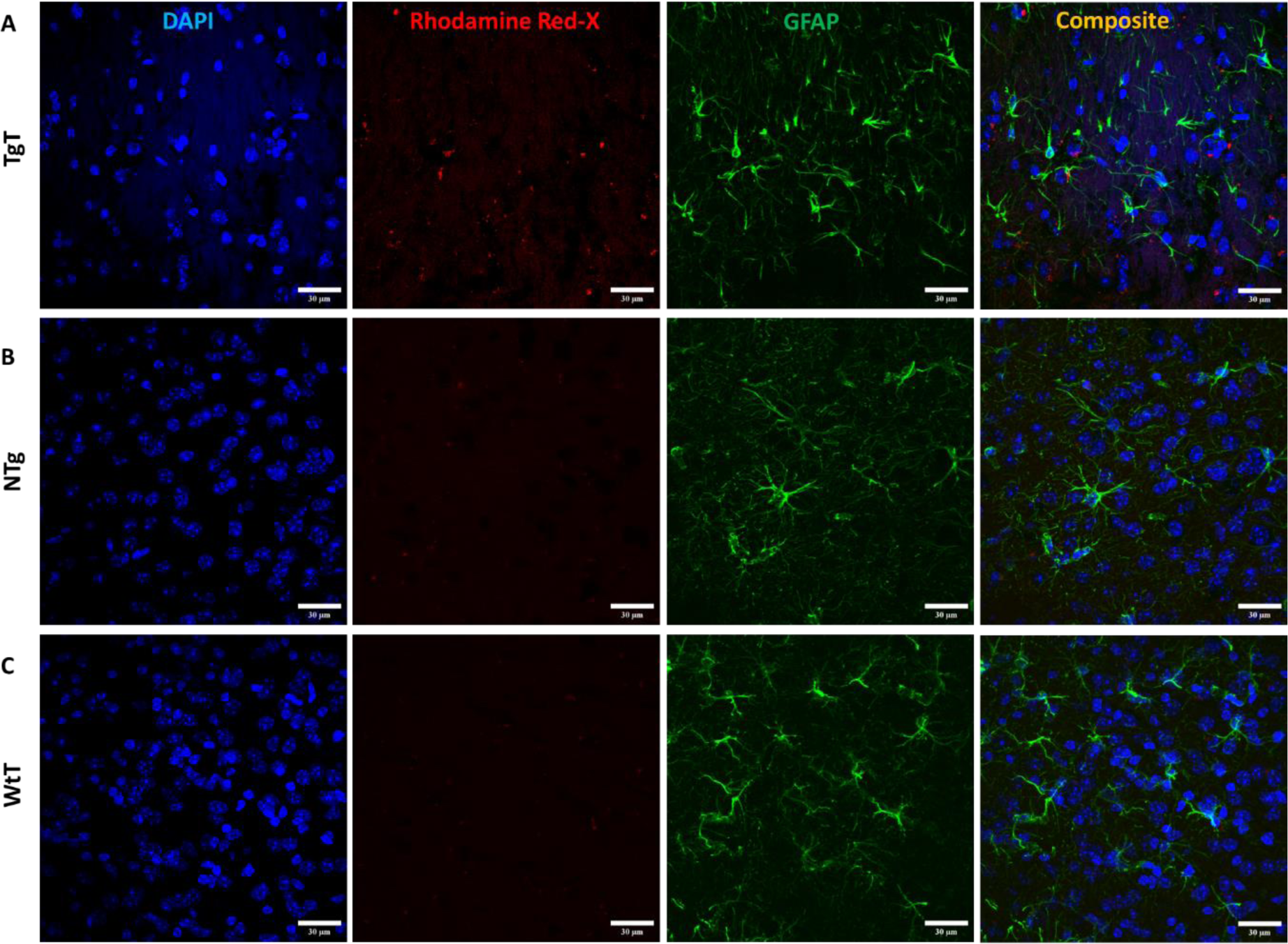
Astrocytes do not display convincing evidence of *in vivo* uptake. **Entorhinal cortex** sections from 13-15-month-old **TgT**, **TgN** and **WtT** mice show cell bodies and processes with strong GFAP reactivity (green fluorescence). **A) TgT** sections show strong NS signal (Rhodamine) but composite images do not show conclusive evidence of internalization of **T** by anti GFAP reactive cells. **B) TgN** and **C) WtT** sections show no signal in the Rhodamine channel. Scale bar, 30 μm.

**Figure S1.19.**
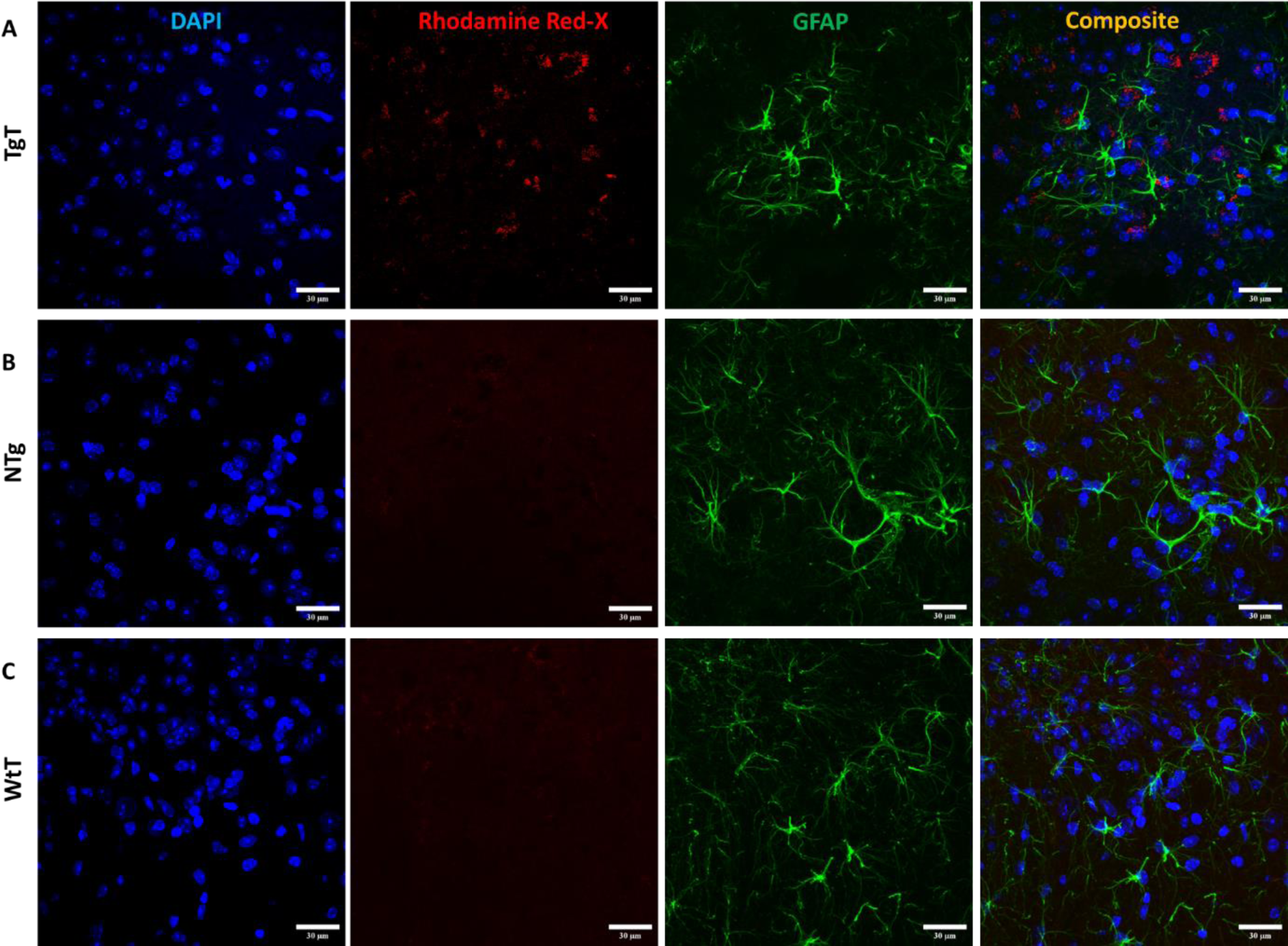
Astrocytes do not display convincing evidence of *in vivo* uptake. **Brainstem** sections from 13-15-month-old **TgT**, **TgN** and **WtT** mice show cell bodies and processes with strong GFAP reactivity (green fluorescence). A) **TgT** sections show strong NS signal (Rhodamine), but composite images do not show conclusive evidence of internalization of **T** by GFAP reactive cells; **B**) **TgN** and **C**) **WtT** sections show no signal in the Rhodamine channel. Scale bar, 30 μm.

**Figure S1.20.**
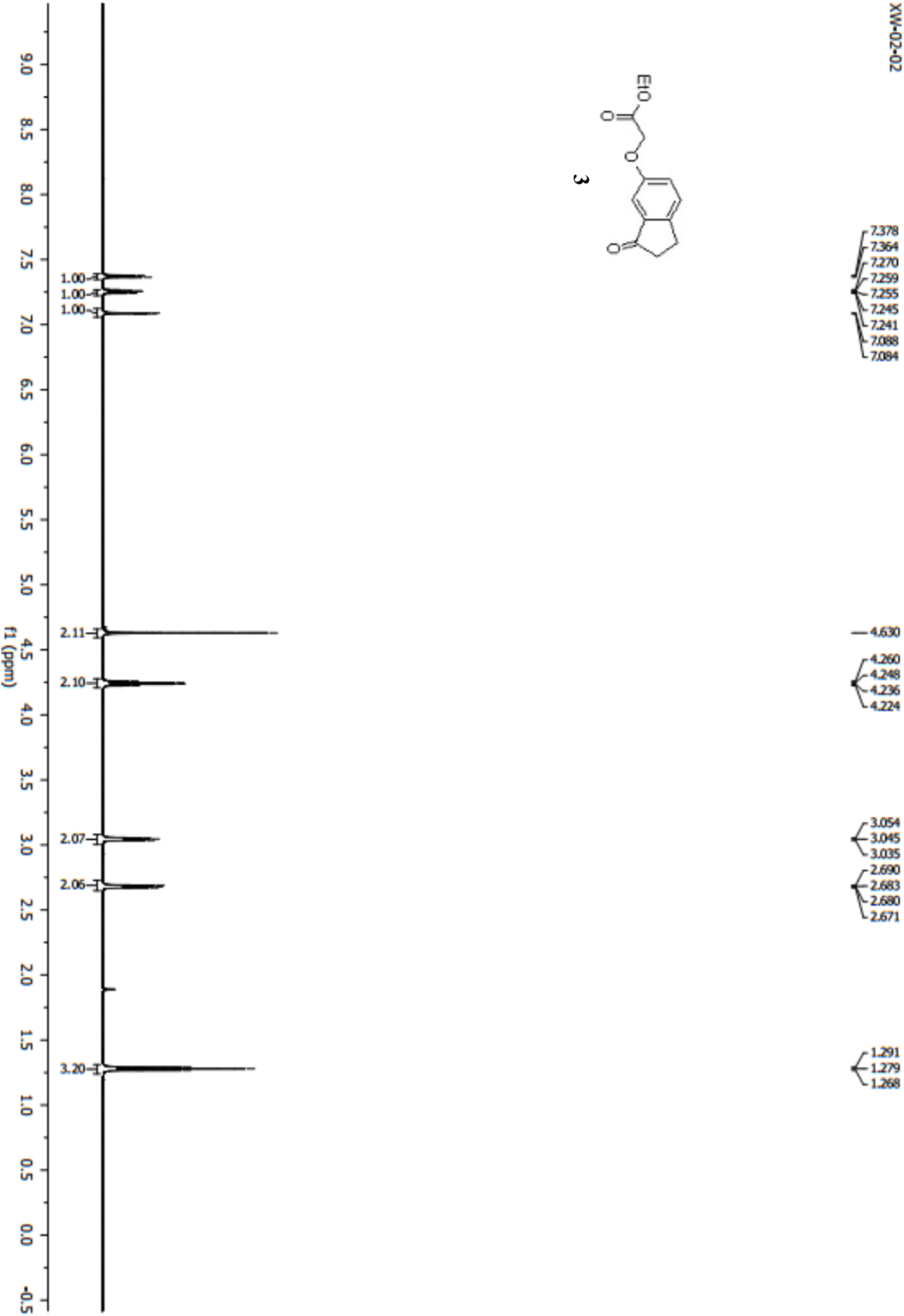
^1^H NMR Spectrum of Compound 3

**Figure S1.21.**
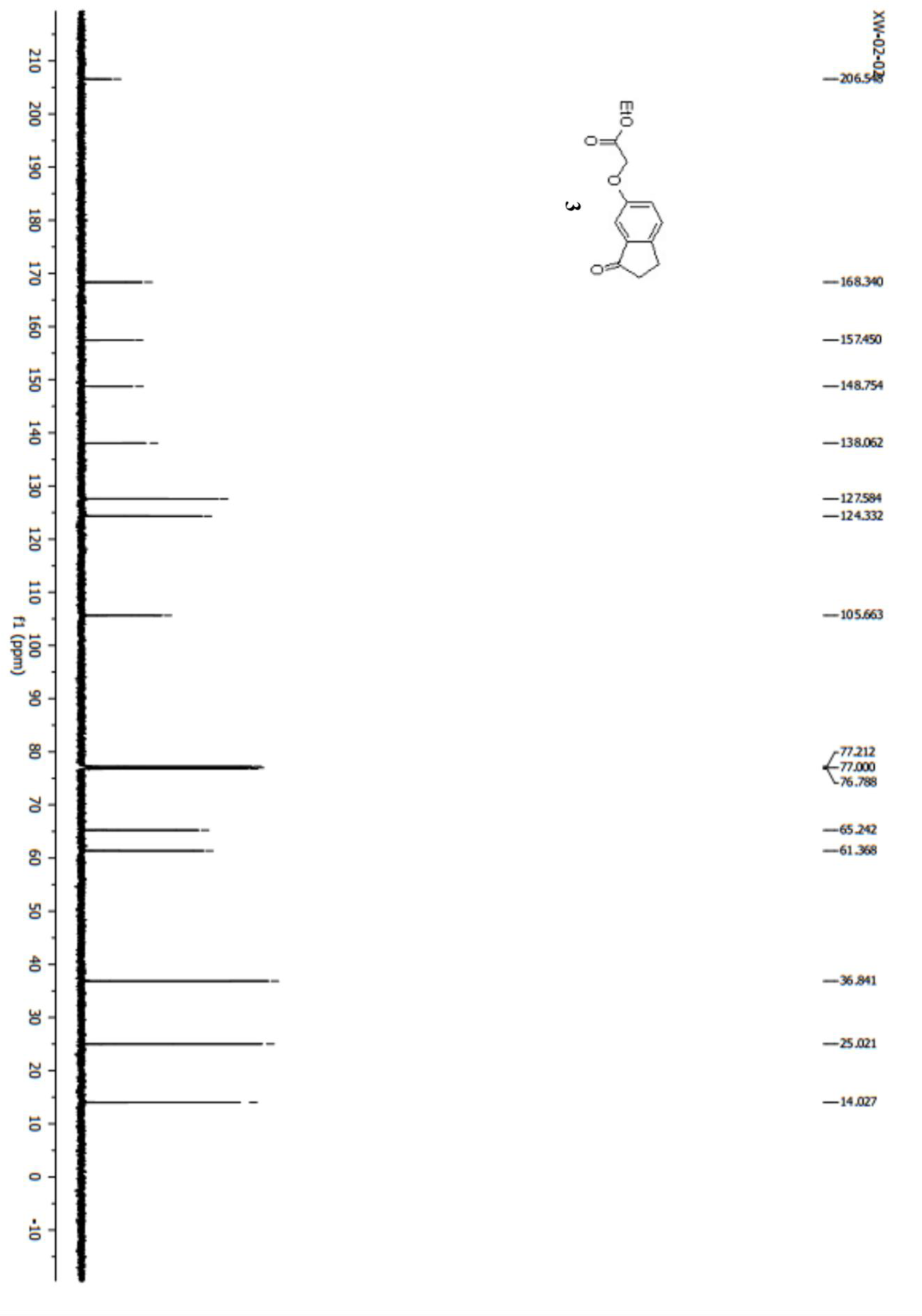
^13^C NMR Spectrum of Compound 3

**Figure S1.22.**
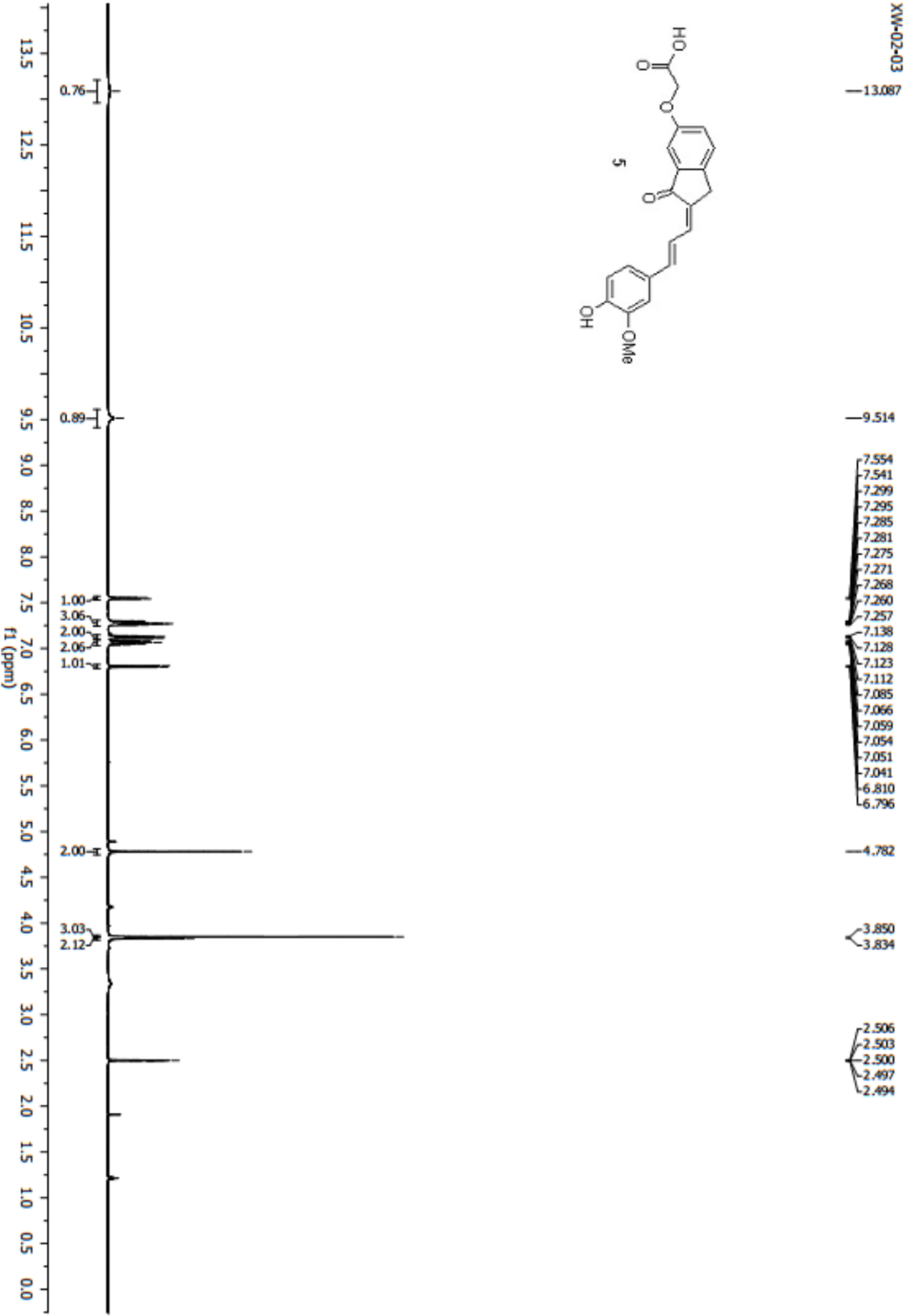
^1^H NMR Spectrum of Compound 5

**Figure S1.23.**
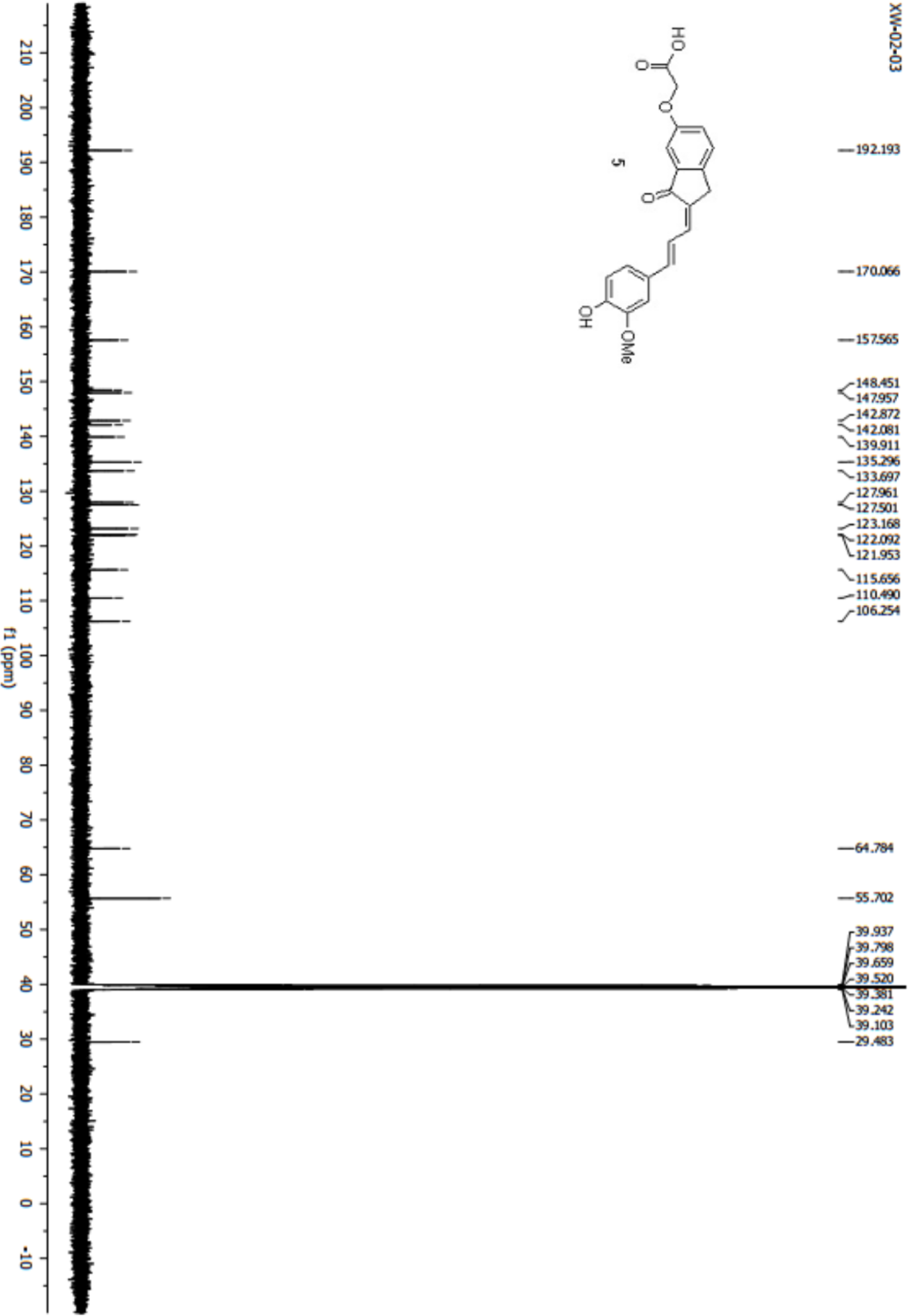
^13^C NMR Spectrum of Compound 5

**Figure S1.24.**
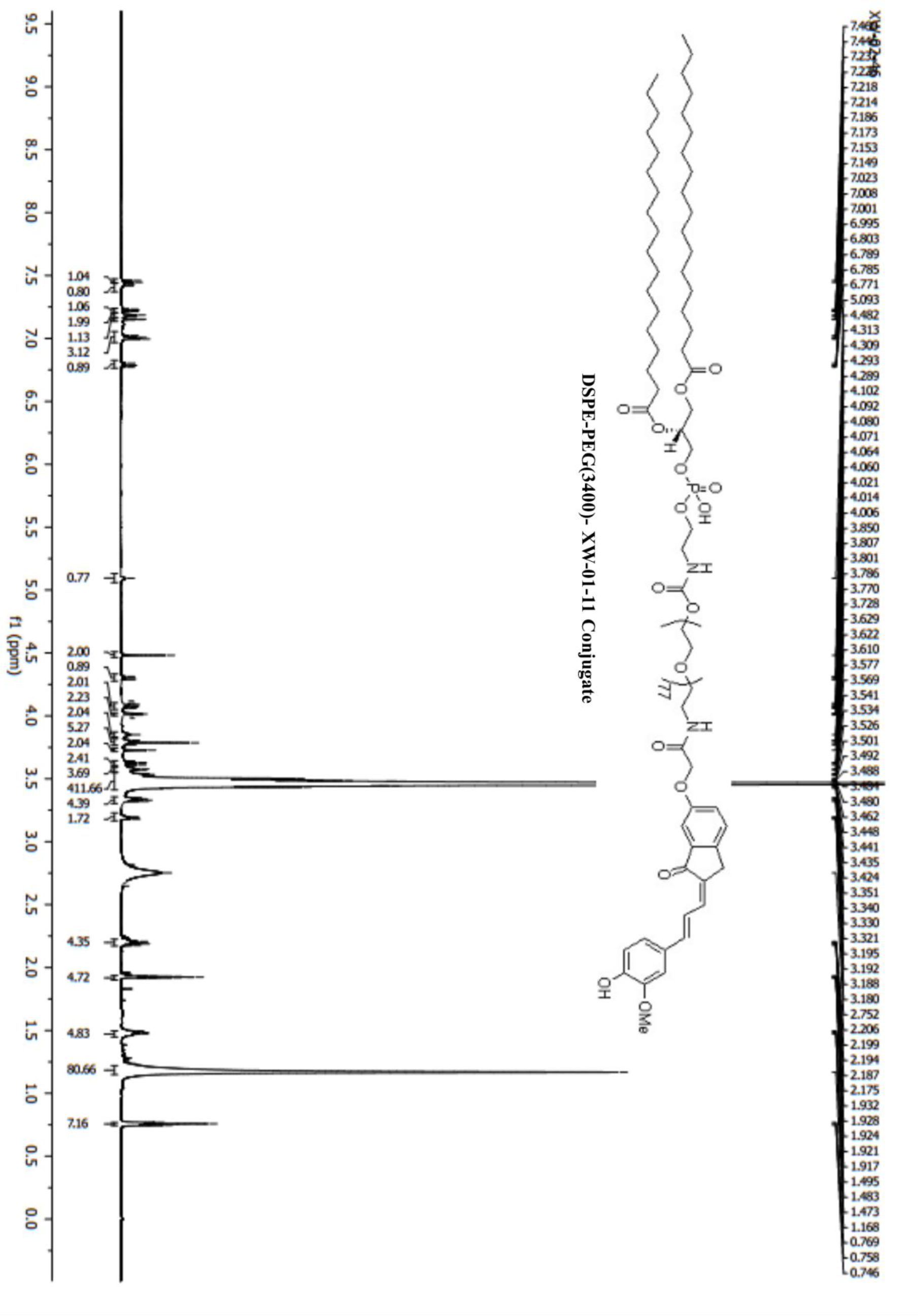
^1^H NMR Spectrum of Compound DSPE-PEG(3400)-XW-01-11 Conjugate

**Figure S1.25.**
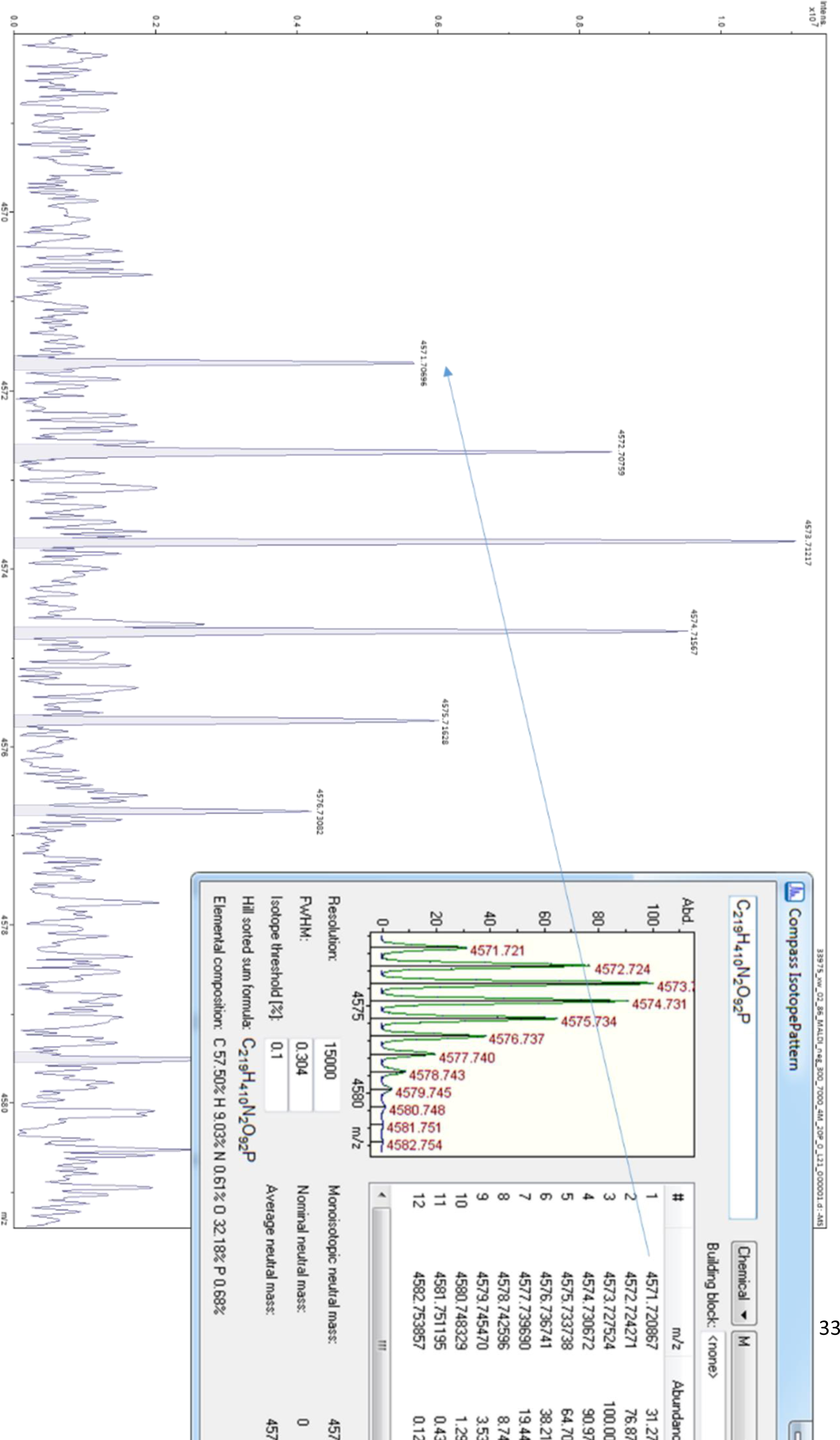
MALDI Spectrum of Compound DSPE-PEG(3400)-XW-01-11 Conjugate

